# Altering cell size asymmetry in *Drosophila* neural stem cells creates supernumerary stem cells with limited lineage expansion potential

**DOI:** 10.1101/2025.09.26.678890

**Authors:** Melissa K Delgado, J Ian Hertzler, Sophia Jannetty, Mia Hoover, Chang Yin, Ilara Yilmaz, Danielle Vahdat, Adam von Barnau Sythoff, Jay Z. Parrish, Neda Bagheri, Clemens Cabernard

## Abstract

Asymmetrically dividing invertebrate and vertebrate stem cells can generate unequal sized sibling cells. However, the functional implications of cell size asymmetry (CSA) are underexplored. Here, we use *Drosophila* neural stem cells (NSCs) to investigate how changes in CSA impact NSC proliferation and cell fate decisions. Using live cell imaging, NSC lineage analysis, and gene expression profiling, we find that altering CSA increases the NSC pool but decreases lineage size and the number of differentiating progeny cells. Modeling CSA *in silico* with a volume-sensitivity and NSC self-inhibition model can recapitulate these findings. Gene expression profiling further revealed that the NSC growth regulator *Imp* and the G1-S cell cycle regulator *CycE* are upregulated in NSCs with altered CSA, providing a potential molecular link to the volume-sensitivity model. We propose that cell size and position regulate NSC proliferation and differential potential, impacting lineage progression and progeny cell differentiation in the developing *Drosophila* brain.

## Introduction

Asymmetric cell division (ACD) is an evolutionarily conserved division mode, generating two daughter cells that acquire different fates ^1^. Binary cell fate outcomes can be achieved via asymmetric segregation of cell fate determinants, including RNA molecules, proteins, or organelles. Alternatively, extrinsic signaling cues can influence cell fate decisions (reviewed in ^2–5^). ACD is used by stem cells to self-renew while forming a differentiating sibling cell at the same time. The correct regulation of ACD is important to prevent tumor formation or neurodevelopmental disorders such as microcephaly ^6–8^.

One manifestation of ACD is the formation of unequally sized sibling cells during mitosis. Sibling cell size asymmetry, referred to as Cell Size Asymmetry (CSA) hereafter, is evolutionarily conserved and has been observed during embryogenesis in ascidians ^9^, leeches ^10^, the roundworm *Ascaris megalocephala* ^11^, sea urchins ^12^, *Ilyanassa obsoleta* (Eastern mudsnail) ^13,14^, *C. elegans* ^15^ and *Drosophila melanogaster* ^16^. CSA also occurs in vertebrate embryos, such as *Xenopus laevis*, where the second division creates an asymmetric embryo with two larger and two smaller blastomeres ^17^. Recent observations further suggest that the first cleavage in the human embryo produces two unequally sized blastomeres ^18^.

Empirical evidence suggests that CSA contributes to cell fate decisions during development. For instance, equalizing the first division in *C. elegans* caused faster cell cycle progression in P1 descendants, defects in cell positioning, division orientation, and cell fate ^15^. In *Volvox carteri*, a multicellular green alga, CSA plays a central role in germ/soma specification ^19^. Changing CSA in asymmetrically dividing *C. elegans* neuroblasts rescued the smaller sibling from apoptosis ^20^. However, how CSA shapes cell fate decisions remain underexplored.

Three main mechanisms have been identified to regulate and implement CSA: (1) spindle displacement through cortical pulling forces, positioning the cleavage furrow machinery off cell center ^7,21,22^, (2) forming and retracting cortical polar lobes into one of the two sibling cells ^13,14,16^, or (3) asymmetric localization of cortical proteins regulating unequal membrane expansion ^2,16,20,23–26^.

*Drosophila* neural stem cells, called neuroblasts (NBs), generate CSA by asymmetrically localizing, and dynamically displacing Actomyosin ^23–25,27–32^. Neuroblasts divide asymmetrically by size and fate, self-renewing the larger neuroblast while either forming a small, differentiating Ganglion Mother Cell (GMC) or an immature Intermediate Progenitor cell (INP). Neuroblasts are intrinsically polarized, manifested in the apical localization of the Par complex, directing the formation of a basal cell fate determinant complex. The latter is composed of the cortical protein Miranda (Mira), the Notch inhibitor Numb (NUMB, NUMBL in humans), the translational repressor Brain Tumor (Brat; TRIM2,3 in humans) and the transcription factor Prospero (Pros; Prox1, 2 in humans). Basal cell fate determinants segregate into the small INP or GMC, while the apical Par complex and associated proteins remain in the self-renewed neuroblast ^33–35^.

In *Drosophila* larval neuroblasts, asymmetric Myosin localization is regulated via intrinsic polarity cues. For instance, in early mitosis, the polarity protein Partner of Inscuteable (Pins; LGN/AGS3 in vertebrates) enriches Myosin on the apical neuroblast cortex. In early anaphase, Pins induces a basally directed cortical flow by recruiting Protein Kinase N (Pkn) to the apical neuroblast cortex, relieving it from Actomyosin contractile tension ^23,28,29,32,34,36^. This release in Actomyosin-dependent cortical tension permits biased membrane expansion to form a large, self-renewed apically positioned neuroblast and a smaller differentiating Ganglion Mother Cell ^24,30,37^.

While the larger NB retains apically localized proteins, Mira, Pros, Brat and Numb segregate into the small GMC promoting the expression of neural differentiation genes and represses neuroblast stem cell identity genes ^38–45^. Although ample experimental evidence supports the model that GMC differentiation is induced through asymmetric segregation of basal cell fate determinants, it is unclear whether also CSA contributes to cell fate decisions and/or stem cell behavior in this system.

Here, we use asymmetrically dividing *Drosophila* neural stem cells to investigate how altered CSA impacts neurogenesis and brain development. We employ two complementary approaches to change CSA of dividing neuroblasts and find that cell positioning and cell size regulates the NBs proliferation and differentiation potential. Surprisingly, live cell imaging, detailed lineage analysis and gene expression profiling shows that changing CSA increases the neural stem cell pool, without increasing the number of differentiating siblings. Using iterative mathematical modeling, we show that volume-based growth regulation and neuroblast self-repression are possible mechanisms to constrain neuroblast lineage expansion. We conclude that the size and placement of the NB is necessary to maintain a proliferative stem cell pool to produce the correct number of differentiating siblings. This finding could inform our current understanding of neurodevelopmental disorders, such as primary microcephaly.

## Results

### Retaining activated Myosin on the apical neuroblast cortex is sufficient to alter CSA

*Drosophila* neuroblasts divide asymmetrically by size, producing a large apically positioned neuroblast and a small, basally located GMC ^35,45^. To understand the role of CSA in *Drosophila* neurodevelopment, we aimed to create developing larval fly brains in which some neuroblasts divide symmetrically by size and/or show inverted physical asymmetry. In the latter case, this should lead to an apically positioned cell that adopts the size of a GMC and a basally positioned cell the size of a neuroblast. To this end, we used a fly strain, harboring the regulatory subunit of Non-muscle Myosin II (Myosin hereafter), encoded by *spaghetti squash (sqh)*, endogenously tagged with EGFP (*Sqh::EGFP*, hereafter) ^46^. Next, we expressed UAS-ALD-RockCA::vhhGFP4 (called nanobody hereafter) with the neuroblast-specific driver *worniu-Gal4* (*worGal4* ^47^) in flies containing Sqh::EGFP. The nanobody construct contains Inscuteable’s apical localization domain (ALD) ^48^ and the constitutively active kinase domain from Rho kinase (RockCA). The apically localized nanobody binds to the EGFP moiety of Sqh::EGFP, tethering and activating it only on the apical neuroblast cortex (Figure S1A) ^29,49^. The apical neuroblast cortex can be identified by the increased concentration of Sqh::EGFP (Figure S1B). Additionally, retaining activated Myosin apically does not disrupt neuroblast polarity ^29^. Consistent with our earlier findings, maintaining a pool of activated Myosin on the apical neuroblast cortex resulted in three division outcomes: (1) approximately 60% normal asymmetric, (2) approximately 27% symmetric, and (3) approximately 8% inverted asymmetric divisions. The inverted divisions produced a small apical and a larger basal cell. Some neuroblasts also displayed a more pronounced NB-GMC cell size ratio; however, these occur less frequently in wild-type and control conditions (Figure S1B, C). Here, we focus on the cellular and developmental consequences of symmetric and inverted asymmetric divisions.

### Sibling cell size and position determines the cell’s division potential

We next asked whether cell size correlates with proliferation potential. Wild-type *Drosophila* neuroblasts are at least twice as large as GMCs and divide every 1 - 4 hours. GMCs have a long cell cycle exceeding 8 hrs ^50,51^. If cell size predicts cell cycle progression, we would expect that small apical cells of inverted neuroblast divisions have a longer cell cycle than their larger basal siblings. Similarly, the siblings resulting from symmetric neuroblast divisions should display similar cell cycle lengths. To test these hypotheses, we performed long-term live cell imaging experiments, imaging control and nanobody-expressing larval brains for 8-14 hrs (see methods). We used the spindle marker mCherry::Jupiter ^51^, expressed with the neuroblast driver *worGal4* ^47^ to assess cell cycle stage and length. As expected, wild-type neuroblasts, expressing Sqh::EGFP in conjunction with mCherry::Jupiter showed cell cycle lengths between 1.5 - 4 hrs and always created a large apical neuroblast and a small basal GMC (Figure 1A, D; Movie 1 and ^51^). Neuroblasts expressing the nanobody in the presence of Sqh::EGFP and mCherry::Jupiter generally divided less frequently (see below). Still, we found occasional inverted or symmetric neuroblast divisions, whereby symmetric divisions occurred more regularly than inverted asymmetric divisions (∼ 73% symmetric, ∼ 27% inverted asymmetric; n = 85; Figure 1B, C, E; Movie 2, 3). Symmetrically dividing neuroblasts, induced through spindle misalignment, divide orthogonal to the neuroblast’s initial apical–basal polarity axis ^52–54^, whereas nanobody-induced symmetric or inverted divisions divide along the apical–basal polarity axis (Figure S1D–G) ^29^. Therefore, nanobody-induced symmetric or inverted divisions maintain the sibling cells’ original position while changing their cell size. We next asked whether the apical or basal cell is more proliferative after nanobody-induced inverted and symmetric divisions. To answer this question, we analyzed lineages of symmetric and inverted neuroblast divisions where either the apical, basal, or both siblings divided again. We found 6 inverted neuroblast divisions where all basal cells divided again. In these lineages, only 2 small apical cells were able to divide within the recorded time frame, but only after the basal cell divided first (Figure 1F, H). Interestingly, in size-symmetric divisions, apical cells were more proliferative than basal cells. For instance, of 20 symmetrically dividing neuroblasts, all apical but only 5 basal cells divided again. Of these 5, 4 apical cells divided before its basal sibling (Figure 1G, I). Our live cell imaging protocols allow to record multiple neuroblast divisions in larval brain explants ^51,55–57^ but although we imaged 10-14 hours, we never detected more than 2 divisions of the apically-positioned cell following a symmetric NB division (Figure 1C). Therefore, nanobody-expressing NBs created short, but asymmetric hemilineages. Based on these results, we conclude that if cell volume is equal between neuroblast siblings, the apically positioned cell is more proliferative. However, if cell volume is unequal, the larger cell is more proliferative.

**Figure 1:**
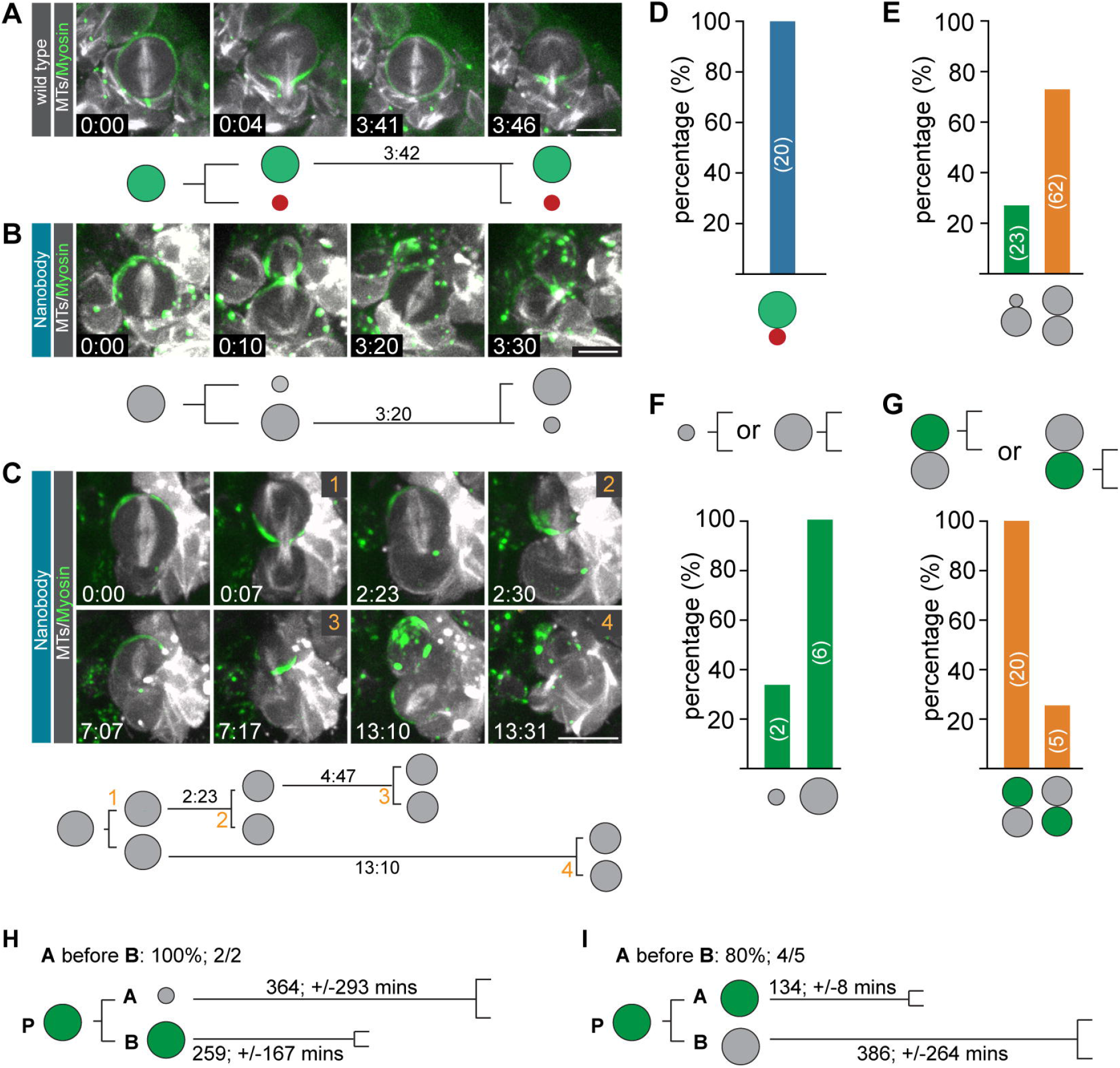
Cell size and position determine the cell’s proliferation potential. Representative image sequence of an **(A)** asymmetrically dividing wild-type, **(B)** nanobody-expressing NB dividing with inverted **(C)** or symmetric cell size asymmetry. All examples show Sqh::EGFP in green and mCherry::Jupiter in white. Division mode of wild-type **(D)** or nanobody-expressing **(E)** neuroblasts. Only inverted and symmetric divisions were counted in nanobody-expressing neuroblasts. **(F)** Quantification of apical and basal cell divisions, following an inverted asymmetric **(F)** or symmetric **(G)** neuroblast division. Quantification of cell cycle time and frequency of apical and basal cell division for **(H)** inverted or **(I)** symmetric neuroblast divisions. Numbers on bars indicate number of observed neuroblasts. Scale bar is 10μm; time in hours:minutes.

### Cell position and cell size determine cell fate outcomes in NB divisions with altered CSA

Next, we asked whether the sibling cells of symmetric or inverted asymmetric neuroblast divisions adopt neuroblast fate. To that end, we used CRISPR/Cas9 to generate a live neuroblast marker by knocking in the fluorescent protein tdTomato ^58^ at *Deadpan’s (Dpn)* endogenous locus ^59^ (see methods). We crossed *tdTomato::Dpn* to *sqh::EGFP* and the *worGal4*-driven nanobody construct. In wild-type neuroblasts, tdTomato::Dpn was localized in the nucleus during interphase and released into the cytoplasm in early mitosis. In telophase, cytoplasmic tdTomato::Dpn segregated into the self-renewed wild-type neuroblast and the differentiating GMC. Shortly after cytokinesis, tdTomato::Dpn was incorporated into the neuroblast’s nucleus but was also visible in the differentiating GMC. However, the neuroblast Dpn signal was stronger compared to the GMC in wild-type neuroblasts (Figure 2A-C; Movie 4). Next, we assayed tdTomato::Dpn segregation and expression in *mud* mutants because symmetrically dividing *mud*^*4*^ null mutant neuroblasts give rise to two Dpn+ sibling cells ^51^. As expected, symmetrically dividing *mud*^*4*^ mutant neuroblasts inherited equal amounts of tdTomato::Dpn, entering the nucleus in both sibling cells. In both equal-sized siblings, tdTomato::Dpn signal was maintained at comparable levels (Figure 2D – F; Movie 5). Similarly, the apical and basal siblings of symmetrically dividing nanobody-expressing neuroblasts inherited and maintained equal amounts of tdTomato::Dpn (Figure 2G-I; Movie 6). Surprisingly, nanobody-expressing neuroblasts dividing with inverted cell-size asymmetry showed two different outcomes: (1) in one instance, we found a small apical cell that showed no detectable tdTomato::Dpn after mitosis (Figure 2J-L; Movie 7). (2) A second event showed a smaller apical cell containing nuclear tdTomato::Dpn (Figure 2M – O; Movie 8). Interestingly, in both instances, the large basal sibling contained tdTomato::Dpn. We conclude that for induced symmetric and inverted divisions, larger sibling cells always receive and maintain nuclear Dpn, while smaller siblings may also inherit and sustain nuclear Dpn.

**Figure 2:**
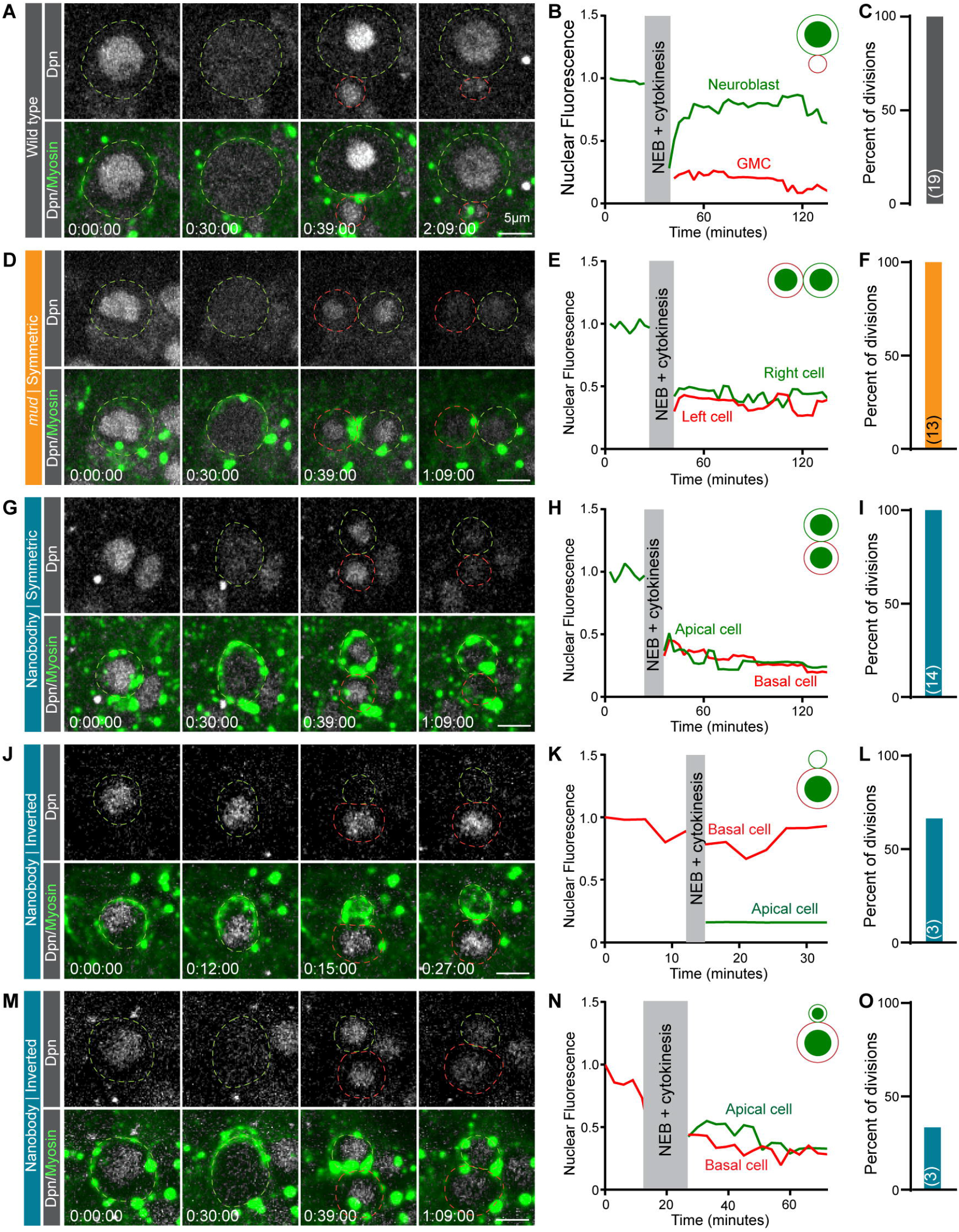
Dpn is inherited by the siblings of symmetrically or inverted asymmetrically dividing neuroblasts. Representative image sequence of a **(A)** wild type, **(D)** *mud*^*4*^ mutant, or **(G, J, M)** nanobody-expressing neuroblasts. All shown examples express tdTomato::Dpn (white) and Sqh::EGFP (green). Dotted lines outline the parental and sibling cells. **(B, E, H, K, N)** Sum of intensity quantification of tdTomato::Dpn over time in the two sibling cells for the shown examples. Schematics in the upper right corner of the graph illustrate the division outcome and Dpn (green) inheritance. **(C, F, I, L, O)** Quantification of division outcomes. Numbers on bars indicate number of observed neuroblasts. Scale bar: 5μm. Time in hours:minutes:seconds

### Larval brains with altered CSA maintain the same cell populations, but with altered ratios

Wild type neuroblasts generate all the cell types in the developing larval brain ^35^. Since changes in CSA alter the number, size, and proliferation potential of larval neuroblasts (see ^51^ and above), we hypothesized that changes in NB CSA could impact the number and type of neuroblast progeny cells. To define the molecular changes and cell fate outcomes of CSA, we sought to investigate gene expression profiles in single cells. To this end, we performed single-cell RNA sequencing on control, *mud*^*4*^, and nanobody-expressing brains at 72 - 96 hours after egg laying (see methods). While control brains only contain asymmetrically dividing neuroblasts, *mud*^*4*^ mutant brains are composed of a mix of asymmetric and symmetrically dividing neuroblasts. Nanobody-expressing brains develop with normal asymmetric, symmetric, and inverted asymmetric divisions (Figure S1D-G). For each condition, we dissected and dissociated approximately 50 larval brains into single-cell suspensions, sorted for cells of interest, and prepared cDNA libraries. We used the standard 10x Genomics Cellranger pipeline for initial alignment, and a Seurat pipeline for quality control, integration, and further analysis (detailed protocols in methods). After quality control, we retained approximately 13,500 cells for each condition. We identified 28 individual clusters and annotated them based on the presence and abundance of validated marker genes, as well as by comparing differentially expressed genes (DEGs) from annotated clusters in published *Drosophila* scRNAseq datasets ^60–64^. We annotated clusters 15 and 19 as neuroblasts based on the expression of *dpn, mira*, and *CycE* among others ^65–70^. Cells in cluster 11 expressed the validated INP genes *ham* and *opa* ^71,72^. Clusters 5, 7, 9, 12 expressed the GMC genes *grh, tap, dap* ^73,74^, clusters 3, 13, 14 the early neuronal marker *Hey* ^75^, clusters 1, 2, 4, 8, 10, 18 the immature neuron markers *elav* and *fne*, but no *nSyb* or *brp* ^76,77^. Several clusters contained cells expressing mature neuronal markers such as *nSyb, brp*, and various neurotransmitter marker genes ^78,79^. We also found more specialized, terminal cell types in our atlas, such as Kenyon cells (*Rgk1*) ^80^, glia (*repo, wrapper*) ^81,82^, and a neuron type with high expression of all three Iroquois gene complex members (*ara, caup*, and *mirr*) ^83^ (Figure 3A, D, and Figure S2-6).

**Figure 3:**
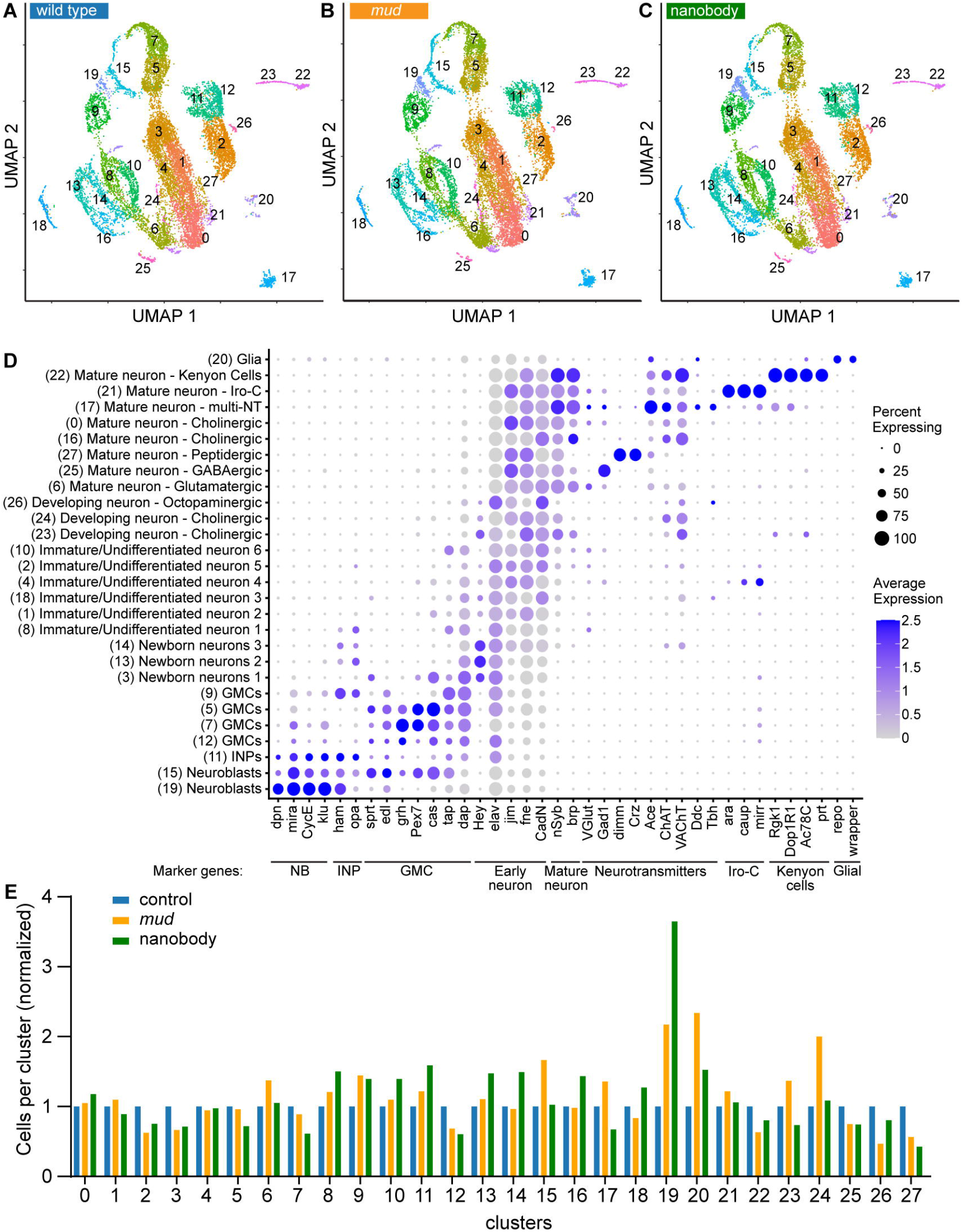
Altering neuroblast sibling cell size asymmetry changes cell numbers but not cell types. UMAP plots of the scRNAseq clusters for **(A)** wild type, **(B)** *mud*^*4*^ and **(C)** nanobody-expressing neuroblasts. All 28 clusters (0-27) are present in all 3 conditions. **(D)** Annotations of all clusters (y-axis) with representative marker genes (x-axis). Dot size indicates the percent of cells per cluster expressing the shown gene. Dot color indicates the relative expression of the gene where zero represents average expression. Values below zero show up as gray, regardless of magnitude. Clusters are ordered vertically by developmental age and type. **(E)** Graph of cell numbers per cluster, normalized for library size, and normalized to control for each individual cluster.

We hypothesized that symmetric and/or inverted asymmetric cell divisions will either (1) create new or hybrid cell types with unique expression profiles, (2) eliminate characterized cell types or, (3) create differences in the number of specific brain cells. Surprisingly, cluster analysis revealed that control, *mud*, and nanobody-expressing brains all contained the same 28 clusters. While the expression profiles of all three genotypes varied (see below), standard clustering methods neither revealed a new, nor prevented the formation of a known cell type compared to the wild-type cell atlas (Figure 3A-D). However, more detailed analysis of cluster sizes showed that certain cell types appeared in increased or decreased numbers (Figure 3E). Most noteworthy, the neuroblast cluster 19 showed the highest difference in neuroblast numbers between the three conditions. These findings suggest that altering CSA in neuroblasts neither creates new, nor eliminates existing brain cell types but changes the number of specific brain cell populations.

### Changes in neuroblast sibling cell size increase the neuroblast pool

In wild-type third instar brains, the neural stem cell pool remains constant, but *mud*^*4*^ mutant brains contain ∼ 50% more neuroblasts, due to symmetric, neuroblast-amplifying divisions ^51,84^. Consistent with this result, the neuroblast clusters 15 and 19 showed more cells in both *mud*^*4*^ mutant and nanobody-expressing brains (Figure 4A, B). To verify this result, and to assay how increasing the neuroblast pool affects neurogenesis, we labelled type I neuroblast lineages in wild type, *mud*^*4*,^ and nanobody-expressing brains with *AseGal4* ^85^, driving the expression of mCherry::Jupiter. Type I wild-type lineages only contain 1 Dpn^+^ neuroblast ^86–88^ and multiple differentiating Pros^+^ cells, providing a clear binary readout. We stained third instar larval control, *mud*^*4*^ mutant, and nanobody expressing brains with anti-Dpn and anti-Prospero. We segmented type I neuroblasts, Pros^+^ cells, and identifiable Ase-Gal4 expressing lineages to quantify cell type, cell size, cell number, and lineage size. While all wild-type type I lineages contained a single Dpn^+^ neuroblast surrounded by multiple smaller Pros^+^ differentiating cells, nanobody-expressing brains often contained two, and *mud*^*4*^ mutant brains up to four Dpn^+^ neuroblasts (Figure 4C-F). Notably, we did not find any nanobody or *mud*^*4*^ mutant lineages with more than four Dpn^+^ neuroblasts. In addition, both *mud*^*4*^ and nanobody-expressing brains contained significantly smaller Dpn^+^ nuclei, indicating that *mud*^*4*^ and nanobody-expressing neuroblasts are smaller than wild-type neuroblasts (Figure 4G). *mud* mutant brains have been reported to contain enlarged mushroom bodies, containing more neuroblasts and differentiated Kenyon cells ^54^. Surprisingly, our segmentation analysis revealed that type I neuroblast lineages are smaller and contain fewer Pros^+^ cells in *mud*^*4*^ and nanobody-expressing brains (Figure 4H, I). While in nanobody-expressing brains the neuroblast’s mitotic index was significantly reduced compared to wild-type brains, *mud*^*4*^ mutant neuroblasts surprisingly showed only a small, albeit significant, decrease in their mitotic index (Figure 4J, K). From these data, we conclude that altering neuroblast CSA increases the neuroblast pool while decreasing neuroblast size. We further conclude that a larger neuroblast pool fails to generate more differentiating cells, thereby restricting lineage expansion.

**Figure 4:**
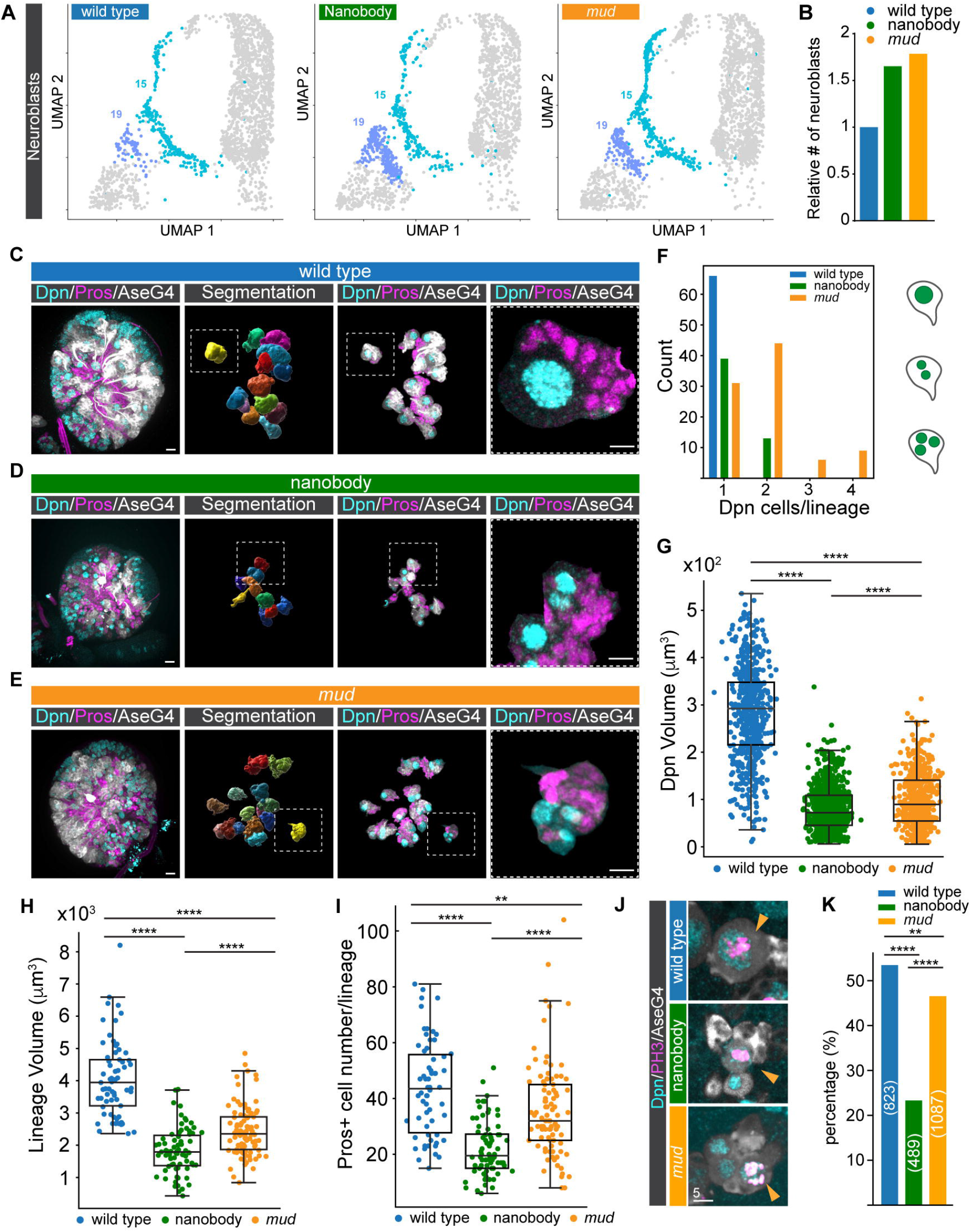
Altering neuroblast sibling cell size asymmetry changes neuroblast size, lineage size and cell number. **(A)** UMAP plot of scRNAseq data for the neuroblast clusters 15 and 19 for each condition. **(B)** Quantification of the relative total neuroblast number, normalized for library size and control neuroblast numbers. Representative type-I (AseGal4 > UAS-mCherry::Jupiter; white) neuroblast lineages for **(C)** wild type, **(D)** nanobody-expressing and **(E)** *mud*^*4*^ mutant neuroblasts stained with anti-Dpn (cyan) and anti-Prospero (magenta). Segmentation of AseGal4-expressing lineages and neuroblasts (not shown) was used to count **(F)** neuroblast number per lineage, **(G)** neuroblast (Dpn) nuclear size, **(H)** lineage size and **(I)** number of Prospero cells per lineage. Representative images **(J)** for wild type, *mud*^*4*^ and nanobody-expressing type I (AseGal4 > UAS-mCherry::Jupiter; white) neuroblasts stained with anti-Dpn (cyan) and anti-PH3 (magenta). **(K)** Quantification of mitotic index. Scale bar is 10μm. p < 0.05 were considered significant; * p < 0.05, ** p < 0.01, *** p < 0.001, **** p < 0.0001. The following statistical tests were used: Anova and Tukey HSD for (G, H, I, K).

### An increased neuroblast pool does not increase the number of GMCs, early-born, immature, or mature neurons

We next wondered whether our RNAseq dataset could be used to determine how changes in CSA affect cell differentiation by assessing the relative numbers of brain cell populations. As a proof of principle, we first compared the number of cells expressing *prospero* mRNA (*pros*^*+*^ cells) with cells immunoractive for the anti-Prospero antibody (Pros^+^ cells). *prospero* is expressed ubiquitously in wild-type, *mud*^*4*^, and nanobody-expressing cells, providing a generic marker for various types of neuroblast progeny (Figure 5A, B). Based on the IHC data, we expected to see a proportional reduction in *pros*^*+*^ cells in our cell atlas in *mud*^*4*^ and nanobody-expressing brains. Indeed, while the fraction of *pros*^*+*^ cells (excluding cluster 15 & 19) is similar across the different conditions (Figure 5A, B), we calculated that one wildtype neuroblast generates approximately 31 *pros*^*+*^ cells based on the RNAseq dataset. This number was reduced in *mud*^*4*^ and nanobody-expressing brains (*mud*^*4*^: 18; nanobody: 20). We next calculated the number of Pros^+^ cells per neuroblast based on our IHC dataset. Consistent with the RNAseq data, we found that wild type neuroblasts generate on average 42 Pros^+^ cells (SD+/-17.7; n = 62 lineages), *mud*^*4*^ mutant neuroblasts produce ∼ 22 Pros^+^ cells (SD+/-14; n = 93 lineages) and nanobody-expressing neuroblasts generate ∼ 20 Pros^+^ cells (SD+/-11; n = 72 lineages) (Figure Figure 5I, J). Thus, the RNAseq and IHC datasets show comparable numbers of Pros+ cells per neuroblast.

**Figure 5:**
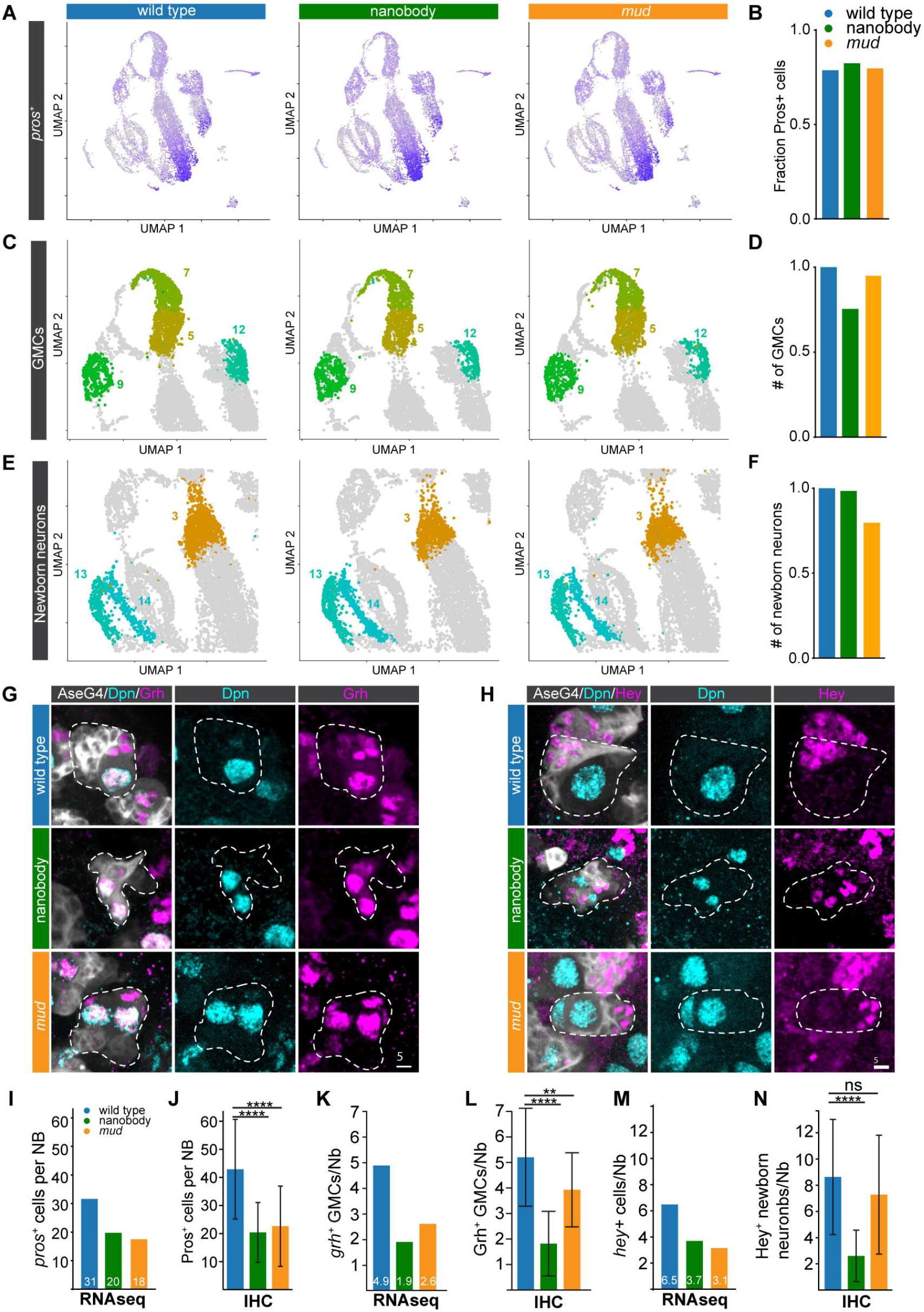
Altering neuroblast sibling cell size asymmetry reduces GMC, newborn neurons and overall prospero-positive cell numbers. **(A)** Plots of *prospero* expression for wild type, nanobody-expressing and *mud*^*4*^ mutant brains overlaid on the UMAP. High Prospero-expressing cells are visualized with intense purple. **(B)** Fraction of cells per condition with at least 1 *prospero* transcript. UMAP plots for wild type, nanobody and *mud*^*4*^ highlighting **(C)** GMC or **(E)** newborn neuron clusters. Quantification of relative **(D)** GMC numbers and **(F)** newborn neurons, respectively, normalized for library size and control. Representative type I neuroblast lineages (AseGal4 > UAS-mCherry::Jupiter; white), stained for **(G)** anti-Dpn (cyan) and anti-Grh (GMCs; magenta) or **(H)** anti-Dpn (cyan) and anti-Hey (newborn neurons; magenta). Quantifications of the number of cells expressing *prospero* (*pros+*) determined by **(I)** RNAseq or **(J)** IHC. Quantifications of the number of cells expressing grainyhead, determined by **(K)** RNAseq or **(L)** IHC. Quantifications of the number of cells expressing Hey determined by **(M)** RNAseq or **(N)** IHC. Scale bars are 5μm. p < 0.05 were considered significant; * p < 0.05, ** p < 0.01, *** p < 0.001, **** p < 0.0001. The following statistical tests were used: One-way Anova and Dunnett’s multiple comparisons test for (J, L, N).

Each asymmetric type I neuroblast division generates a GMC, and we wondered whether an increase in the neuroblast pool also increases the number of GMCs. The transcription factor Grainyhead (grh) ^89^ is a specific marker for the GMC clusters 5, 7, 12 (Figure 3D, Figure S3C, D). Per RNAseq data, a wild-type neuroblast should generate ∼ 5 *grh*^*+*^ GMCs, whereas *mud*^*4*^ mutant and nanobody-expressing NBs are expected to produce approximately 3 and 2, respectively (Figure 5K). To validate these numbers, we stained wild-type, *mud*^*4*^, and nanobody-expressing brains with anti-Dpn and anti-Grh antibodies and counted the number of Grh^+^ cells per neuroblast. Consistent with the RNAseq results, we calculated 5.2 Grh^+^ cells per neuroblast in wild type (SD+/-1.9; n = 59 lineages), 3.9 Grh^+^ cells in *mud*^*4*^ (SD+/-1.4; n = 28 lineages) and 1.8 Grh^+^ cells in nanobody-expressing neuroblasts (SD+/-1.3; n = 70 lineages) (Figure 5G, K, L). We also assessed the number of newborn neurons, using Hey as a marker (Figure 3D and Figure S4A, B). We again found similar numbers of Hey^+^ cells per wild-type neuroblast comparing RNAseq and IHC data, and both *mud*^*4*^ and nanobody-expressing brains had a significant reduction in Hey^+^ cells (Figure 5H, M, N). Finally, our RNAseq dataset also highlighted a decrease in *elav*^*+*^ immature and *nSyb/brp*^*+*^ mature neurons in *mud*^*4*^ and nanobody-expressing brains (Figure S6B-G). We conclude that despite an increase in the neuroblast pool, the number of GMCs, early-born, immature, or mature neurons decreases in larval type I NB lineages with altered CSA.

### Modeling predicts that unregulated CSA would exceed NB, GMC and neuron numbers

Symmetrically dividing *mud*^*4*^ mutant neuroblasts have a reduced mitotic index (Figure 4K) but still divide frequently compared to wild type NBs ^51^. Thus, it is surprising that the increase in the neural stem cell pool does not exceed 4 NBs per lineage, which fail to generate more differentiating progeny than observed in wild type lineages. We used an ordinary differential equation (ODE) model to better understand the consequences of neuroblast division mode on cell differentiation and cell numbers. We first developed a base model that describes three core processes of NB lineage progression: NB division, GMC division, and neuron maturation. The cell types in the model include NBs, GMCs, immature neurons, and mature neurons (Figure 6A). In the base model, NBs divide either symmetrically, yielding two NBs, or asymmetrically, yielding one NB and one GMC. Model parameters include NB and GMC division rates, neuronal maturation rate, and the fraction of NB divisions that are symmetric. To simulate wild type lineage development, we initialized the model with one neuroblast, set the fraction of symmetric divisions to 0%, and parameterized the division and maturation rates from experimental data (see methods). The wild-type simulation produced 1 NB, 6 GMCs, and 35 immature neurons, comparable to what we observe *in vivo* (Figure S7A).

**Figure 6:**
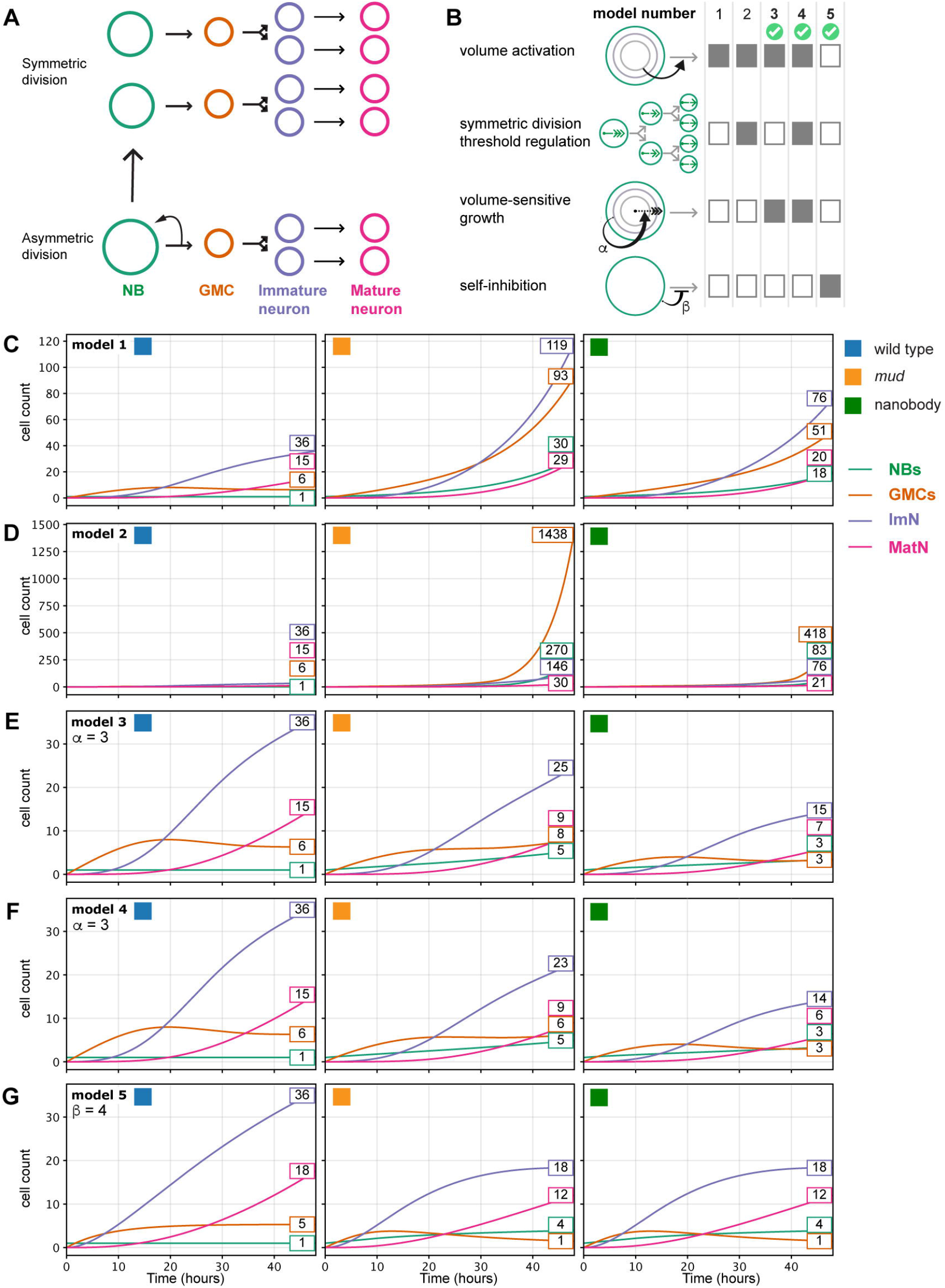
ODE modeling shows volume-sensitive NB growth and NB self-inhibition are both sufficient mechanisms for constraining NB lineage expansion. **(A)** Cartoon showing the differentiation cascade of NBs following asymmetric (bottom) and symmetric (top) division. **(B)** Summary of dynamics included in each model. Bold numbers and green check marks denote models in which mutant proliferation was effectively constrained. **(C-F)** Cell counts for each model over the duration of the simulation for wild type (left), *mud*^*4*^ mutant (center), and nanobody (right) simulations.

To simulate mutant NB lineage development, we increased the fraction of symmetric divisions to 15%, consistent with the rate of symmetric divisions experimentally observed in *mud*^*4*^ mutant NBs ^51^. This simulation yielded a lineage with a substantially higher cell count than the wild-type simulation (122 NBs, 326 GMCs, 596 immature neurons; Figure S7B). This mismatch between observed cell counts *in vivo* and *in silico* model output suggests that additional regulatory mechanisms restrict the NB pool size and the number of differentiating siblings after symmetric divisions.

### Volume-dependent division dynamics are insufficient to explain mutant lineage behavior

Our base model does not account for cell growth or volume, but it is known that NBs regrow after each division during in third instar larval brains ^35,90^. To evaluate whether growth- and volume-dependent regulatory mechanisms could explain the reduced size of mutant lineages, we extended the base model to explicitly track both cell counts and cell volume (Figure 6A, B). In this framework, there is one volume term for each cell type that tracks the total volume of all cells for the given cell type. Simulations are initialized with one NB with an initial volume of 905µM^3^. NBs grow at a constant parameterized rate and divide either asymmetrically, or symmetrically. In this model, when NBs divide asymmetrically, the NB volume term retain 80% of the dividing NB volume and pass 20% to the GMC volume term. Asymmetric divisions increase the GMC count by one, whereas the NB count remains unchanged (1 NB total). When NBs divide symmetrically, the NB volume term retains the full dividing NB volume and the NB count increases by one. GMCs also grow and divide in our simulations, passing their volume to neurons; neurons do not grow or divide.

To regulate NB division, we implemented a volume-activation mechanism (Figure 6B) in which division rates follow a power-law dependence on average NB size. In this model, the NB’s growth rate is constant, but their division rates are very low until their volume approaches a threshold, set to 125% of their initial volume. At that threshold the division rate increases sharply. A lower bound is imposed so NBs with volumes less than 25% their initial size are unable to divide.

We first asked whether this volume-activation mechanism was sufficient on its own to constrain mutant proliferation. Because NBs produced by symmetric divisions begin smaller and NB growth rate is constant between mutant and and wild type NBs, we expected the mutant NBs to take longer to reach the division threshold, prolonging their cell cycles and potentially suppressing mutant NB lineage proliferation. To test this theory, we simulated *mud* mutant NB colonies that had 15% symmetric divisions, and nanobody-expressing NB colonies that had both 15% symmetric divisions and an NB growth rate reduced to 80% of wild type. As expected, *mud* and nanobody-expressing NBs showed prolonged cell-cycle durations relative to wild type (Figure S8A, B). However, this delay was insufficient to offset the expansion of the NB pool, and mutant simulations still predicted substantial lineage overgrowth (Figure 6C; Model 1).

Experimental evidence indicates that *mud*^*4*^ mutant NBs generated through symmetric division do not regrow to their full initial volume but divide at smaller volumes than wild type NBs (Figure S9A, B). To capture this observation, we introduced an additional dynamic in which the NB division volume threshold decreased incrementally with each symmetric division (Figure 6B; Model 2). As expected, this modification accelerated proliferation in mutant simulations, ultimately driving *mud* mutant and nanobody models to the maximum NB division rate, causing an increase in cell counts that is inconsistent with experimental observations (Figure 6D; Figure S8C).

Together these results show that despite the smaller volumes of NBs resulting from symmetric divisions, volume-based division alone is insufficient to constrain mutant NB lineages. Thus, additional negative feedback must act on NB growth and/or division to suppress symmetrically dividing NB overgrowth.

### Volume-sensitive growth could suppress excessive lineage expansion of symmetrically dividing NBs if volume sensitivity is high

The preceding sections established that some form of negative regulation must act on symmetrically dividing neuroblasts (NBs) to prevent excessive lineage expansion and NB pool amplification. A salient phenotypic difference between NBs produced by symmetric versus asymmetric divisions is starting size. As noted above, symmetrically produced NBs are smaller than the pre-symmetric division parental NB (Figure S9A, B). In contrast to model 1 and 2, where NB growth rate was constant across all NBs, we decided to evaluate whether volume-sensitive growth would effectively suppress mutant NB proliferation. In this model, smaller NBs grow at a slower rate than larger NBs: when the average NB volume was smaller than a reference value (set to equal the initial NB volume), the growth rate was reduced below the base rate. As the average NB volume exceeded the reference, the growth rate increased proportionally. We introduced a sensitivity parameter, *α*, to control how sharply the growth rate responds to volume. Low *α* values cause the NB growth rate to rise gradually as the average NB volume increases; high *α* values strongly suppress growth when average volume is below the reference and accelerate it once the reference is exceeded.

We implemented this volume-sensitive growth in two settings: (i) with a static volume threshold for division (Model 3), and (ii) with a threshold that decreases with each symmetric division (Model 4) (Figure 6B). Our simulations show that volume-dependent growth can suppress mutant proliferation if growth-volume coupling is sufficiently strong. For *α* ≤ 2 both models predicted that symmetrically dividing mutant NBs outgrew wild-type lineages. For *α* ≥ 3, the slowing of growth in smaller NBs was sufficient to reduce lineage cell counts in *mud* and nanobody-expressing NBs compared to wild type. Notably, Model 4 consistently predicted fewer cells than Model 3 at a given *α* (Figure 6E, F; Figure S8D, E; Figure S10, Figure S11). Although lowering the division threshold with each symmetric in Model 4 decreases the amount the NBs must grow to divide, by allowing cells to divide at smaller volumes it also allows NB starting size to decrease. At high alpha values, where NB growth is strongly volume-sensitive, this reduction feeds back to further suppress NB growth, ultimately extending the cell cycle duration and constraining proliferation more strongly than in Model 3 (Figure S8D, E). Consistent with these modeling results, we measured cell cycle durations in *mud*^*4*^ mutant neuroblasts that are similar to what model 3 and 4 predicted (Figure S9C). Similarly, nanobody-expressing NBs, dividing symmetrically or inverted asymmetrically generate siblings with extended cell cycle times and reduced proliferation rates (Figure 1C, H, I).

Taken together, Models 3 and 4 suggest that strong coupling of NB growth rate to NB volume is sufficient to restrict NB numbers and lineage expansion after symmetric NB divisions. We conclude that volume-sensitive growth is a possible regulatory mechanism if NB growth is highly sensitive to cell size.

### Neuroblast self-repression can suppress lineage expansion of symmetrically dividing NBs

We next asked whether the size of the neuroblast pool itself could act as a regulatory signal. Specifically, we wondered whether a negative feedback mechanism, in which neuroblasts repress their own division and/or growth could effectively constrain NB lineage expansion (Figure 6B; model 5). To test this, we extended the base model to include a Hill-type negative feedback mechanism, repressing NB division rates. Hill-type feedback mechanisms represent regulatory processes in which the strength of the (in this case, inhibitory) regulation increases as the regulator abundance increases. In our system, an increase in NB number per lineage would increase the inhibition on the NB’s division rate. The inhibition strength is regulated by the Hill coefficient: a low coefficient increases repression gradually as a function of NB number, while higher coefficients (here, *β*>=2) strongly inhibit division once the NB count passes 2. Our simulations showed that for *β*>=3, NB self-repression was sufficient to reduce *mud* and nanobody-expressing NB lineage sizes below wild-type cell counts (Figure 6G; Figure S8F; Figure S12). Together these results show that NB self-repression can effectively suppress lineage expansion in *mud* and nanobody-expressing NBs.

### *mud*^*4*^ and nanobody-expressing neuroblasts differentially express neuroblast and differentiation genes

Both our experimental and modeling data highlight that lineage expansion is mainly driven by NBs proliferation behavior. *mud*^*4*^ and nanobody-expressing neuroblasts are sufficiently similar to wild type NBs to form the same clusters. Nevertheless, we aimed to use gene expression to better define their molecular identity. We performed a differential expression analysis of cluster 15 and 19, separately comparing gene expression profiles of *mud*^*4*^ and nanobody-expressing neuroblasts with wild-type neuroblasts. To ensure we retained the most significant Differentially Expressed Genes (DEGs), we only considered genes that are expressed in at least 25% of neuroblasts, had a p-value of 0.001 or better, and a minimum log2-fold change of 0.5. This left us with 161 DEGs between wild type and *mud*^*4*^, and 104 between wild type and nanobody (Figure 7A-D). In our initial analysis, we noticed several genes associated with neuroblast or neuronal cell fate that were differentially expressed in *mud*^*4*^ or nanobody-expressing neuroblasts compared to wild-type NBs (highlighted in Figure 7A, C). For instance, the neuroblast markers *miranda* (Mira) was upregulated in *mud*^*4*^ and nanobody-expressing neuroblasts. *Nervous finger 1 (nerfin-1)*, a transcription factor required for the maintenance of larval neuron differentiation ^91,92^, or *ham*, a PRDM family transcription factor controlling intermediate precursor cell maturation in the type II larval neuroblast lineages ^93^, were also differentially expressed in both *mud*^*4*^ mutant and nanobody-expressing neuroblasts (Figure 7F). To group our DEGs into ‘NB’ and ‘differentiation’ genes, we checked whether the identified DEGs are preferentially expressed in NBs or neurons, using a published dataset as a reference ^94^. This analysis revealed that genes differentially expressed in both *mud*^*4*^ and nanobody-expressing NBs were slightly enriched for genes that are normally expressed in wild type NBs (Figure 7F).

**Figure 7:**
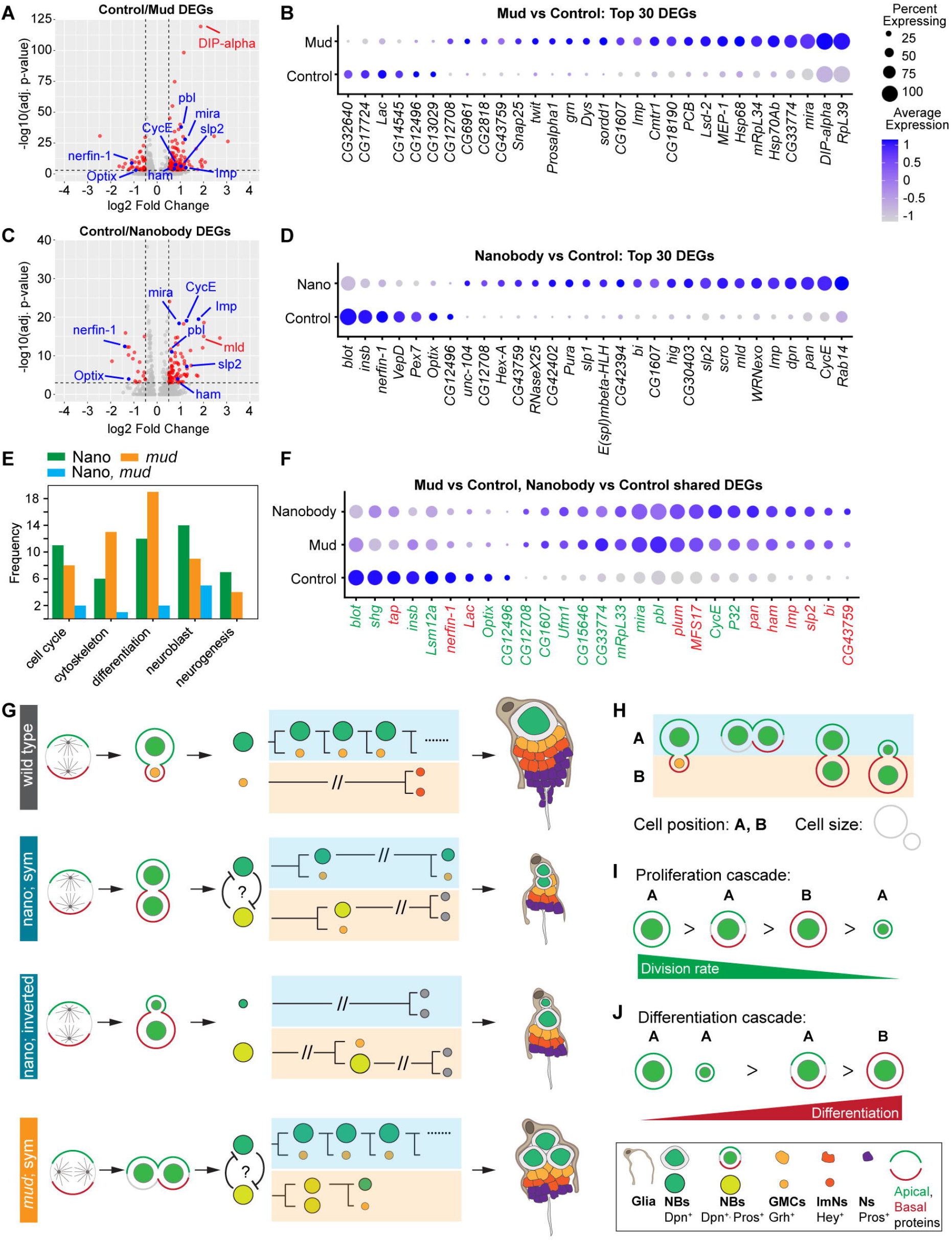
*mud*^*4*^ and nanobody-expressing neuroblasts differentially express genes associated with neuroblast fate, the cell cycle, cell differentiation, neurogenesis, and the cytoskeleton. Volcano plots of differentially expressed genes (DEGs) comparing **(A)** wild type with *mud*^*4*^, and **(C)** wild type with nanobody-expressing neuroblasts, respectively. DEGs with an adjusted p-value of 0.001 or lower and a log2FC of 0.5 or higher are plotted with colored dots. Annotated dots refer to genes of interest. Dotplots of the top 30 DEGs comparing **(B)** wild type with *mud*^*4*^, **(D)** wild type with nanobody-expressing neuroblasts, and **(F)** DEGs shared between *mud*^*4*^ and nanobody-expressing NBs compared to wild type, respectively. Genes annotated in green and red correspond to genes expressed in NBs and neurons, respectively ^94^. Dotplots only show the top 30 DEGs by log2FC. **(E)** Frequency of selected keywords associated with the DEGs. **(G - J)** Summary and Model. NBs dividing with altered CSA (symmetric or inverted asymmetric) create shortened and asymmetric lineage trees, resulting in diminished lineage expansion (different cell types are indicated with colors, see legend). **(H)** The position and size of sibling NBs (A, B), generated with altered CSA define their **(I)** proliferation rates and **(J)** differentiation potential.

We next performed a GO term analysis and searched FlyBase for specific keywords to get functional insight into the DEGs. Interestingly, while several GO terms highlighted neuroblast fate commitment, cell proliferation, or neurogenesis processes, we found many genes associated with the keywords ‘cell cycle’, ‘cytoskeleton’, ‘differentiation’, ‘neuroblast’ or ‘neurogenesis’ (Figure 7E and Figure S13 & Figure S14). In conclusion, *mud*^*4*^ mutant and nanobody-expressing neuroblasts differentially express genes that are associated with neuroblast expression or function, as well as genes associated with differentiation, cell cycle or cytoskeleton.

### The NB size regulator *Imp* and cell cycle regulator *CycE* are both upregulated in *mud* and nanobody-expressing neuroblasts

After this initial characterization, we screened our DEGs for genes and pathways involved in NB growth to find a potential mechanism for the proposed volume-based coupling to cell cycle progression. Wild type NBs exiting quiescence activate cell growth and cell cycle entry via the Pi3K/Akt pathway in response to secreted Insulin-like peptides via the Insulin Receptor (InR) ^95^. Similarly, the progressive reduction in neuroblast cell size in pupal stages is mediated by Ecdysone signaling, the mediator complex, and oxidative phosphorylation ^50^. The p21-activated kinase Mushroom bodies tiny (Mbt; PAK4 in vertebrates) and the Casein Kinase 2 target Mushroom body miniature (Mbm) are also implicated in NB growth and proliferation control ^96,97^. Except for *Mld*, a transcription factor required for production of the steroid hormone ecdysone ^98^, none of these pathways showed differentially expressed genes in either *mud*^*4*^ or nanobody-expressing brains (Figure S15A-D). However, the mRNA-binding protein *Imp* (IGF2BP1 in vertebrates), regulating NB size and division rates ^99^, is upregulated in *mud*^*4*^ and nanobody-expressing NBs (Figure 7A - D, F). Interestingly, *mud*^*4*^ mutant neuroblasts also upregulated *DIP-alpha*, a member of the immunoglobulin superfamily (IgSF) cell adhesion molecules. DIP-alpha has been shown to localize to NB membranes, interacting with Dpr10 and Dpr6, which are expressed in cortex glia. This pathway has been proposed to regulate NB proliferation via mechanical signaling between NBs and cortex glia ^100^. Strikingly, both *mud*^*4*^ and nanobody-expressing brains upregulated the cell cycle regulator *CycE*. CycE regulates entry into S-phase and overexpression of CycE accelerates cell proliferation without stimulating cell growth ^101^. CycE also regulates cell fate in embryonic NBs, a function that is independent of its role in the cell cycle ^102^. We conclude that while most genes from the Insulin, mediator complex and oxidative phosphorylation pathway are not differentially expressed, the neuroblast growth gene *Imp* and the cell cycle regulator and neuroblast marker *CycE* are both upregulated in *mud*^*4*^ and nanobody-expressing neuroblasts.

### Dpn and Pros targets are differentially regulated in *mud* and nanobody-expressing NBs

We finally searched our DEGs for clues of how NB self-inhibition could be implemented molecularly. In wild type NBs, GMC differentiation and thus NB inhibition is induced via asymmetric segregation of basal cell fate determinants, such as Prospero, Numb or Brat ^35^. Miranda serves as a shuttle for the transcriptional repressor Prospero ^35,45^. The basal cell fate determinant Mira can still segregate asymmetrically in symmetrically dividing *mud*^*4*^ or nanobody-expressing NBs ^23,29,51^. This is consistent with our live cell imaging data: equal-sized *mud*^*4*^ mutant, originating from symmetric divisions, show different proliferation behavior, reflected in the fact that one sibling divides more frequently that the other. Similarly, basal cells of symmetrically dividing nanobody-expressing NBs are not as proliferative as their apical siblings. These differences in sibling NB proliferation rates gives rise to asymmetric hemilineages (Figure 1F-I; Figure S9B).

We wondered whether *mud*^*4*^ and nanobody-expressing NBs would differentially express Pros targets. We found several DEGs shared between *mud*^*4*^ and nanobody-expressing brains that are differentially expressed in *pros* mutants ^42^. For instance, the Pros targets *CycE, bi, mRpL33, mira* and *pbl*, were upregulated, whereas *Lac* and *blot* were downregulated in *mud*^*4*^ *and* nanobody-expressing neuroblasts (Figure S15D).

We also considered the fact that Dpn, a repressor of cell differentiation in the nervous system ^103^, segregates symmetrically in *mud*^*4*^ and nanobody-expressing neuroblasts (Figure 2D-I). Thus, we hypothesized whether Dpn could induce neuroblast-fate by repressing cell differentiation genes. We screened our DEGs, shared between *mud*^*4*^ and nanobody-expressing brains for Dpn-target genes ^103^ and found that *CG15646, ham, imp, mira, slp2* are up-, whereas *nerfin-1* is downregulated in *mud*^*4*^ and nanobody-expressing genes (Figure S15E). In conclusion, *mud*^*4*^ and nanobody-expressing neuroblasts differentially express genes downstream of Pros and Dpn transcriptional regulators. This suggests that Pros and Dpn segregation in symmetrically, or inverted asymmetrically dividing neuroblasts impacts their gene expression profile.

## Discussion

Cell Size Asymmetry is an evolutionary conserved form of Asymmetric cell division. However, how sibling cell size differences relate to cellular function and cell fate decisions is mostly unknown. Here, we used *mud* mutants and an apically localized nanobody to alter sibling cell size asymmetry in *Drosophila* neural stem cells. *mud* mutant neuroblasts can divide symmetrically by size due to spindle orientation defects ^51^. In contrast, nanobody-mediated maintenance of activated Myosin on the apical neuroblast cortex in mitosis induces either inverted asymmetric or symmetric neuroblast divisions ^29^. Importantly, neither *mud* nor nanobody-expressing neuroblasts compromise neuroblast polarization ^29,51,52^. However, *mud* mutants divide orthogonal to the apical-basal polarity axis while nanobody-expressing neuroblasts divide in parallel to the intrinsic polarity axis ^29,52–54^.

Here, while both *mud* and nanobody-expressing brains increase the neuroblast pool in the developing brain, neuroblasts are smaller under these conditions compared to wild type. Live cell imaging data also showed that symmetrically and inverted asymmetrically dividing neural stem cells segregate Dpn into both siblings, giving rise to two equally sized Dpn^+^, or one smaller and one larger Dpn^+^ cell. Our live cell imaging data further revealed that neuroblasts created from inverted or symmetric divisions show strong size and position-dependent proliferation behavior. For instance, we only found two instances where the small, apically positioned neuroblast siblings divided again. And in these instances, larger basal siblings divided before the apical cell. In contrast, apically positioned neuroblast siblings originating from symmetric divisions are more proliferative than their equal sized basal siblings. Siblings of symmetrically dividing *mud* mutant neuroblasts also showed asymmetric proliferation behavior, whereby one sibling divided more frequently than the other (Figure 7G).

Surprisingly, we discovered that despite the increase in neural stem cells, differentiating cell types do not increase in number. Both *mud* and nanobody-expressing brains have smaller neuroblast lineages, with fewer GMCs, immature and mature neurons. We used mathematical modeling to find mechanisms that could recapitulate the observed cell numbers and found that either a volume-sensitive mechanism, restricting neuroblast proliferation when neuroblast volumes are reduced, or a direct NB self-inhibition mechanism could recapitulate the observed cell type numbers.

Gene expression profiling further shows that *mud* and nanobody-expressing neuroblasts upregulate genes associated with neuroblast fate or neuronal differentiation, but also genes implicated in cytoskeletal functions, the cell cycle, or neurogenesis.

Based on these data, we propose that neuroblast size and position regulates cell cycle progression and lineage expansion (Figure 7G-J). Larger, apically positioned neuroblasts are more proliferative, producing more differentiating siblings than smaller apical or equally sized basal siblings. These differences in proliferation create hemilineage asymmetry, whereby one sibling generates more differentiating progeny than its sibling. In both instances, the neuroblasts’ proliferation ability could be regulated by asymmetrically segregating cell fate determinants such as Miranda or Pros. In fact, we previously showed that symmetrically dividing *mud* mutants still segregate Miranda asymmetrically ^23,51^. Since apical Myosin sequestration does not change the NB’s intrinsic polarity, basal siblings most likely still inherit Miranda and Prospero in neuroblasts expressing the nanobody construct. Wild-type GMCs inherit Prospero, which induces the expression of differentiation genes, while downregulating neural stem cell and cell cycle genes ^42^. However, since Dpn segregates symmetrically, entering the nucleus in symmetrically, or inverted asymmetrically dividing *mud* and nanobody-expressing neuroblasts, Prospero-induced differentiation could be offset by Dpn, which is known to repress differentiation genes ^103^. Neuroblasts in *mud* and nanobody-expressing brains also upregulate genes associated with neuroblast fate, such as *mira* or *CycE*. CycE has a dual role in cell cycle progression and neuroblast fate decisions ^102^. Overall, symmetrically, and inverted asymmetrically dividing neuroblasts generate siblings that retain neural stem cell fate, but with reduced proliferation potential, which creates asymmetric neuroblast lineages, preventing normal lineage expansion.

All the cells in the developing wild type *Drosophila* brain originate from asymmetrically dividing neuroblasts ^35^. *mud* mutants contain supernumerary mushroom body neuroblasts and increased numbers of Kenyon cells ^54,104,105^. This suggests that in contrast to the type I neuroblasts we investigated here, mushroom body neuroblasts lack certain regulatory mechanisms to rein in the production of extra neurons. Based on our modeling, type I neuroblasts could use volume-sensitivity or NB self-inhibition as extra levels of regulation. While we ultimately do not know the molecular nature of the volume-sensitive and NB self-inhibition mechanisms, it is noteworthy that we find *Imp* and *CycE* upregulated in both *mud* and nanobody-expressing brains. While Imp is required for neuroblast growth, CycE drives cell cycle progression. Similarly, changes in CSA create NBs in direct contact with each other, and cell-cell adhesion-based mechanisms could regulate their cell proliferation behavior. Several genes upregulated in either *mud* or nanobody-expressing neuroblasts encode for cell adhesion molecules (e.g *plum, lac, shg*) or members of the Notch signaling pathway, such as E(spl)mgamma-HLH, or E(spl)beta-HLH. Interestingly, we found the Immunoglobulin superfamily member DIP-alpha upregulated in *mud* mutant neuroblasts. DIP-alpha has been proposed to regulate NB proliferation in *Drosophila* brains via NB-cortex glia interactions. Thus, it is possible that increasing the neural stem cell pool in this cellular niche could provide a regulatory mechanism to regulate stem cell proliferation potential. It is noteworthy that equal sized siblings in *mud*^*4*^ and the apical sibling in nanobody-expressing NBs would be in contact with cortex glia, possibly regulating their proliferation behavior (Figure 7G).

In vertebrate systems, stem cells often undergo an amplification step to increase the size of the stem cell pool, which is necessary to produce differentiating progeny in sufficient numbers ^106–109^. Primary microcephaly, a neurodevelopmental disorder manifested in small brains and reduced cognitive functions, has been proposed to be a consequence of altered neural stem cell divisions: a premature shift towards asymmetric neural stem cell divisions could result in an insufficiently expanded neural stem cell pool and thus a reduction in the number of neurons ^110,111^. However, here we show that an increase in the neural stem cell pool leads to a reduction in neuronal numbers, suggesting that additional mechanisms need to be considered to understand the developmental consequences of altered CSA under physiological normal and disease conditions.

## Methods

### Experimental model and subject details

The following alleles and deficiencies were used: *mud*^*4* 104^. Transgenes and fluorescent markers: *worGal4, UAS-Cherry::Jupiter, Sqh::GFP* (2^nd^ chromosome) ^23^, *worGal4, UAS-Cherry::Jupiter, Sqh::GFP* (3^rd^ chromosome) ^28^, *worGal4, UAS-mCherry::Jupiter* ^51^, *Sqh::EGFP* ^46^, *tdTomato::Dpn* (this work). *pUAST-attB-ALD-Rock CA::VhhGFP4::HA* ^29^, *Ase-Gal4* ^112^. The following recombinant chromosomes were generated using standard genetic procedures: *mud*^*4*^, *sqh::EGFP*.

### Immunohistochemistry

Primary antibodies used in this study were rat anti-dpn (1:100, Abcam 11D1BC7), mouse anti-pros (1:500, gift from Chris Doe), rabbit anti-phospho-Histone-H3 (1:3000, Sigma Aldrich, H0412), rabbit anti-Hey (1:100, gift from Chris Doe), rabbit anti-grh (1:500, gift from Melissa Harrison).

Secondary antibodies used were goat anti-rat (AlexaFluor 647, Invitrogen A21247), goat anti-mouse (AlexaFluor 405, Invitrogen A31553), goat anti-rabbit (AlexaFluor 405, Invitrogen A31556), goat anti-rabbit (AlexaFluor 647, Invitrogen A21245). All secondary antibodies were used at 1:1000. All experiments were conducted with at least 1 overnight in primary and 1 overnight in secondary antibody.

Larvae were dissected by removing the posterior half and inverting, then removing organs, and severing the connections between the back end of the brain and the mouth parts, to minimize stretching of the brain during fixation. Dissected larvae were put in schneiders insect media until fixation. Fixation was performed in 4% PFA in schneiders medium on a nutator for 20 minutes. PFA was then washed off with PBSBT (PBS, 1% BSA, 0.1% Triton-X) 3 times for 20 minutes on a nutator at room temperature. Antibodies were diluted in PBSBT and samples were incubated on a nutator at 4C. After primary staining, samples were washed with PBSBT 3 times for 20 minutes on a nutator at room temperature. Secondary antibody, also diluted in PBSBT, was then applied and incubated on a nutator at 4C. After secondary staining, samples were again washed 3 times for 20 minutes on a nutator at room temperature. Brains were then dissected off of the cuticles and put into vectashield. Brains were mounted on glass slides in a small channel of vectashield between two spacers (glass coverslips), with a third coverslip on top. All samples were imaged immediately after mounting on slides.

### Generation of tdTomato::Dpn CRISPR line

First, tdTomato was PCR amplified from pTIGER_tdTomato::3xFLAG::dCrk (Plasmid #131138, Addgene) with the following primers:

5’ GCGGCCGCGGACATATGATGGTGAGCAAGGGCGAGG 3’

5’ TATACGAAGTTATAGGTACCCTTGTACAGCTCGTCCATGCC 3’

The destination vector, pHDR-SSPB-eGFP-DsRed ^46^, was cut with KpnI and NdeI to remove the SSPB-eGFP, and tdTomato was inserted in its place upstream of LoxP sites (which flank a DsRed for visual confirmation of successful transformation). We then generated a genomic prep of BDSC#58492 (injection stock) and sequenced across the *deadpan* gene region to confirm left and right homology arm (LHA and RHA) sequences with primers:

5’ GTCCGATTCCGATCCATTGC 3’

5’ AGATGGTCGGGCTAATTGTTTG 3’

We used FlyCRISPR target finder tool (http://targetfinder.flycrispr.neuro.brown.edu/) to find CRISPR/Cas9 cut sites. LHA and RHA were amplified by PCR from genomic DNA extracted from BDSC#58492 using the following primers:

LHA: 5’ GAAGCAGGTGGAATTCCAAGTAAAAAGTAAAAAAAAGT 3’

5’ TTGCTCACCATCATATGTTTGTAACGATTTTAATTTGTATATGCAAGTG 3’ RHA:

5’ AGTTATACAGGCGCGCCATGGATTACAAAAACGAT 3’

5’ ACGGAAGAGCCTCGAGGTCGCTCGCCAGCCCCGC 3’

The tdTomato destination vector was digested with EcoRI, NdeI, AscI, XhoI, to excise tdTomato and open the vector backbone. Homology arms with 15bp sequence homology were subsequently inserted into the linearized backbone together with the recovered tdTomato DNA using In-Fusion Cloning (Takara BioSciences).

For the guide RNA, we digested pU6-BbsI-chiRNA ^113^ (#45946, Addgene) with BbsI, and inserted the guides by annealing the two overhanging oligos:

5’ GTAATACGACTCACTATAGGGC 3’

5’ AAACAGTGGGAGATAATCCAAGATC 3’

The finished plasmids were sent to Bestgene Inc (Chino Hills, CA) for injection into embryos. Transformants were delivered and screened for eye DsRed expression. The DsRed cassette was floxed out by crossing to hsCre.

### Live cell imaging

Brains were dissected from animals with fine forceps (Dumont #5, Electron Microscopy Sciences, item number 0103-5-PO) in Schneiders insect media with 10% bovine growth serum (HyClone, item number SH30541.03) and placed in 8-well slides (Ibidi, catalogue no. 80826). Brains were imaged with an Intelligent Imaging Innovations (3i) spinning disk confocal system, with a Yokogawa CSU-W1 spinning disk unit and two Prime 95B Scientific CMOS cameras, and a 60x/1.4NA objective. Imaging was done for 3 -16 hours with the lowest possible laser power and exposure time to minimize bleaching.

### Cell dissociation

10mg/ml, 50ul aliquots of collagenase and papain were made upon receipt of the enzyme with Hank’s Balanced Salt Solution as the solvent, and kept at -20C. Rinaldini solution (100ml milliQ water, 800 mg NaCl, 20 mg KCl, 5 mg NaH2PO4, 100 mg NaHCO3, 100 mg Glucose) was prepared and kept at room temperature. Dissociation solution (400ul Rinaldini buffer, 50ul collagenase, 50ul papain) as well as PBS+1%BSA were prepared on ice immediately before dissection, and kept on ice for the duration of the experiment. All tubes used in this experiment were DNA Lo-Bind tubes. PBS+1%BSA was made fresh and sterile-filtered twice before use. First, larvae were washed briefly in room-temperature PBS before dissection to remove fly media particles. We dissected ∼50-60 brains for each condition in PBS+1%BSA. Approximately 10 brains were dissected before removing the media and replacing with fresh media. All brains were immediately put in an Eppendorf tube on ice containing 500ul PBS+1%BSA. We dissected for one hour before moving on to the dissociation procedure. As much media as possible was removed from the tubes, and then the brains were washed once with cold Rinaldini buffer. Then, 500ul of dissociation solution was added to each tube and tubes were placed at 30C for 30 minutes on a thermomixer with 1000rpm of agitation. Tubes were removed every ten minutes (3 total times) and brains were manually triturated 100x with a P1000 pipette. This cell solution was then filtered through a 40um filter into ice cold FACS tubes. Finally, 0.5ul of far red live/dead cell stain was added (Thermo Fisher L34973). These cell solutions were transported to the FACS sorter on ice. Besides the 30 minutes in the thermomixer, samples and tubes were kept on ice for the full duration of the experiment.

### FACS sorting

Cells were sorted on a BD FACSAria 3 sorter with a 100um nozzle and a flow rate of 1.2. The threshold rate was around 1000-1300 events/second. We targeted 60,000 sorted events per condition, and it took approximately 10-15 minutes of sorting to meet or exceed this number. Cells were sorted for GFP+, mCherry+, and Live/Dead-fluorescence. All cells were sorted into fresh DNA LoBind tubes containing cold 500ul PBS+1%BSA. After sorting all three genotypes, tubes were centrifuged at 300g for 10 minutes at 4C. Excess supernatant was removed, leaving about 60ul of liquid in which to resuspend cells. Cell concentration was noted to be around 600-700 cells per microliter.

### Single cell RNA sequencing

To prepare our cDNA library, we used the 10x Genomics Chromium GEM-X 3’ V4 kit, targeting 20,000 cells per condition. The protocol we followed for this kit can be found at (https://cdn.10xgenomics.com/image/upload/v1710230393/support-documents/CG000731_ChromiumGEM-X_SingleCell3_ReagentKits_v4_UserGuide_RevA.pdf). We performed GEM generation and RT incubation right after dissection and sorting, put the samples at 4C overnight, and finished the cleanup, cDNA amplification, and library construction and sent the samples for sequencing the following day. Sequencing was done by MedGenome Inc (now Signios Biosciences, Foster City, CA) on a NovaSeq X+ with 1 10B lane, and 1.25B paired (2.5B total) reads split across libraries. Samples were demultiplexed by MedGenome Inc and fastq files were delivered.

### scRNAseq Analysis

Samples were initially processed with the CellRanger (10x Genomics) Count function with a custom genome. The base Drosophila Melanogaster genome was downloaded from Ensembl (release 112), and sequences for GFP and mCherry were added to the.gtf and.fa files. All subsequent analysis was done in R. Initial QC was performed for each genotype separately: empty droplets were filtered out with DropletUtils, and low quality cells were removed (<0.8 log10(Genes/Counts), total UMIs <1000, total genes <500, mitochondrial DNA content >18%, ribosomal DNA content <5% or >40%, heat shock genes >5%). Doublets were predicted and removed with the DoubletFinder package. All three genotypes were then integrated into a single Seurat object. PCA analysis was run and the first 30 PCs were used. Clustering was done at a resolution of 1.0, which yielded 28 clusters. Annotation was performed by comparing expression level and abundance of validated marker genes, as well as other DEGs for annotated clusters from other datasets. Dotplots, DEGs, and other expression analyses were performed in R with Seurat.

### Cluster annotation

Clusters were annotated based on validated marker expression. Neuroblasts had high expression of validated markers *dpn, mira*, and *CycE*, and non-validated but consistent markers *Mcm7, Mcm3, PCNA*, and *klu*. INPs had high expression of validated markers ham and *opa*, though little of validated marker *erm* ^72^. GMCs, while all slightly different, shared expression of validated markers *tap* and *dap*, and non-validated but consistent markers *edl, grh, Pex7*, and *insb*. Newborn neurons were characterized by high expression of *Hey* ^75^. Immature/undifferentiated neurons lacked *Hey, nSyb*, and *brp*, but contained varying levels of neuronal genes *elav, fne*, and *CadN* ^76,77^. Mature neurons were characterized by expression of *nSyb* and *brp*, as well as a neurotransmitter marker, such as *Gad1* and *VGAT* for GABA-ergic cells ^114^, *VGlut* for glutamatergic cells ^115^, *VAChT* and *ChAT* for cholinergic cells ^116^, *dimm* and *Crz* for peptidergic cells ^117^, and *Tbh* for octopaminergic cells ^118^. Our Kenyon cell cluster had high expression of many consistent markers, such as *Rgk1, jdp, prt, Dop1R1, Ac78C*, and *PKA-R1* ^80,119^. The glial cluster was the only one that contained any *repo* ^81^. Therefore, our cell atlas represents cell types from neuroblast to differentiated neuron (Figure 5A, B and Figures S2-5).

### Image analysis and quantification

#### tdTomato::Dpn

To quantify tdTomato::Dpn fluorescence over time, we used ImageJ to make a max intensity projection for the range that the nucleus in each neuroblast and GMC occupies, then quantified sum of the pixel intensities (called rawintden in ImageJ) within the nuclei over the duration of the video. We used background subtraction to correct for microscope minimum values. After background subtraction, numbers were normalized to 1 (value of nucleus of first frame shown in representative image sequence).

#### IHC image segmentation and analysis

to quantify Dpn volume, lineage volume, Pros+ cell number and Dpn number/lineage, we used Imaris 9.6 (and higher) to segment Dpn^+^ and Pros^+^ nuclei. Similarly, lineages were segmented and manually corrected if necessary. For lineage volume quantification, axonal projections were manually cut to account for image depth variations in our datasets. Segmentation was done by training Imaris’ pixel classifier or by using threshold segmentation. Grh^+^ and Hey^+^ cells were counted in Imaris using spot detection and counting.

Cell diameter and lineage analysis was performed by manually measuring NB diameters and averaging over three measurements. Cell cycle length was extracted from Imaris. All lineage, diameter, cell cycle, volume and cell count measurements were analyzed in Jupyter notebooks with custom Python scripts.

### GO term network and keyword analysis

For the GO term network and keyword analysis, we only considered DEGs that were expressed in at least 25% of neuroblasts, had a p-value of 0.001 or better, and a minimum log2-fold change of 0.5. DEGs were then converted to dataframes in Jupyter notebooks and analyzed with GSEApy’s Enrichr (GO_Biological_Process_2018).

Keyword analysis was performed with custom Python code in Jupyter notebooks.

## Statistical analysis

Statistical analysis was performed in either Prism or in Jupyter notebooks.

## Code

Code used for RNAseq analysis was written in R. Code used for image, GO-term, and keyword analysis was written in Jupyter notebooks/Python. All the code used in the manuscript is available on github:

## Software and figures

Images and figure panels were assembled and annotated in Adobe Illustrator 2023 (and higher). Image contrast and brightness was adjusted for better visibility, without changing the underlying data.

### RNA sequencing data availability

RNA sequencing data has been submitted to NCBI GEO. Accession number: **GSE308236**.

Seurat files and code can be found on github:

### ODE Modeling

All models were written and executed in Python. Code is publicly available at this URL: https://github.com/Jannetty/Neuroblast-Lineage-ODE-Model/blob/main/ode_model_manuscript_figures.ipynb. All model equations and parameters can be found in jupyter notebook and all simulations can be regenerated accordingly. All models are initialized with one NB. In models that track volume the initial volume of the NB is set to 905µM^3^, a value based on our experimental observations.

#### Base Model

The Base Model tracks cell counts of each cell type in a developing NB lineage.

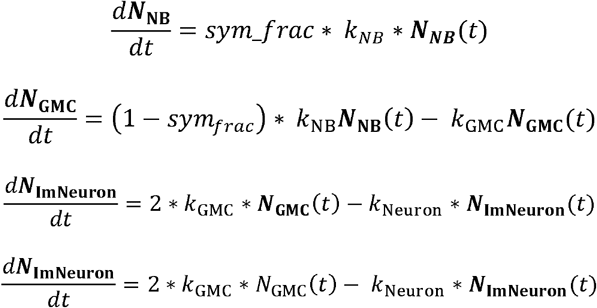

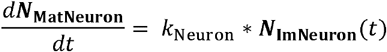

Where ***N***_**NB**_ represents the number of cells in the NB population, ***N***_**GMC**_ represents the number of cells in the GMC population, ***N***_**lmNeuron**_ represents the number of cells in the neuron population, and ***N***_**MatNeuron**_ represents the number of cells in the mature neuron lineage.

The parameter values are as follows:

| Symbol | Parameter | Value | Units | Justification |
| --- | --- | --- | --- | --- |
| sym_frac | Fraction of NB divisions that are symmetric (all others are asymmetric) | 0 for WT<br>.15 for mutants | Unitless | Experimental observation |
| $k_{\text{NB}}$ | Rate of NB division | .67 | $\frac{1}{\text{hr}}$ | Experimental observation |
| $k_{\text{GMC}}$ | Rate of GMC division | .11 | $\frac{1}{\text{hr}}$ | Experimental observation |
| $k_{\text{Neuron}}$ | Rate of neuron maturation | .021 | $\frac{1}{\text{hr}}$ | Experimental observation |

#### Model 1

Model 1 includes the equations and parameters from the Base Model and adds the following

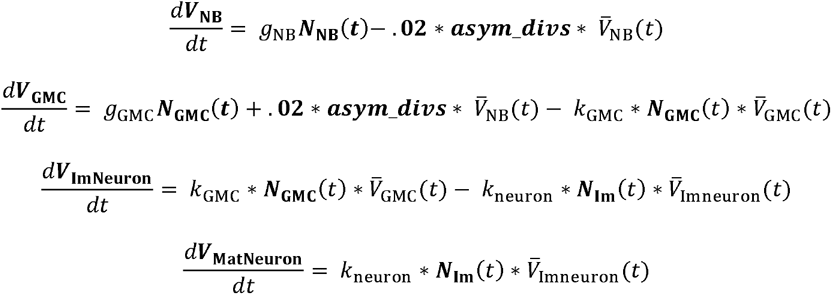

Where *V*_*_ is the total volume of all clels of that cell type in the simulation 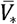 is the average volume of that cell type’s cells 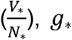 is the per-cell growth rate of the cell type, and *asym*_*divs* is the number of asymmetric divisions, equal to (1 − *sym*_ *frac*) *k*_NB_ ***N***_**NB**_. The NB and GMC division rates are now defined by:

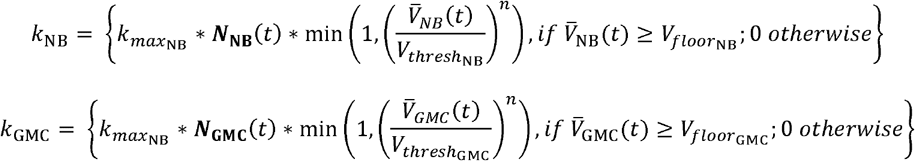

Where 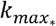 is the maximum growth rate of the cell type, 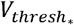 is the division volume threshold for the cell type, 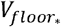 is the minimum volume a cell of the cell type must reach before dividing, and *n* is the sensitivity coefficient.

The new parameter values are as follows:

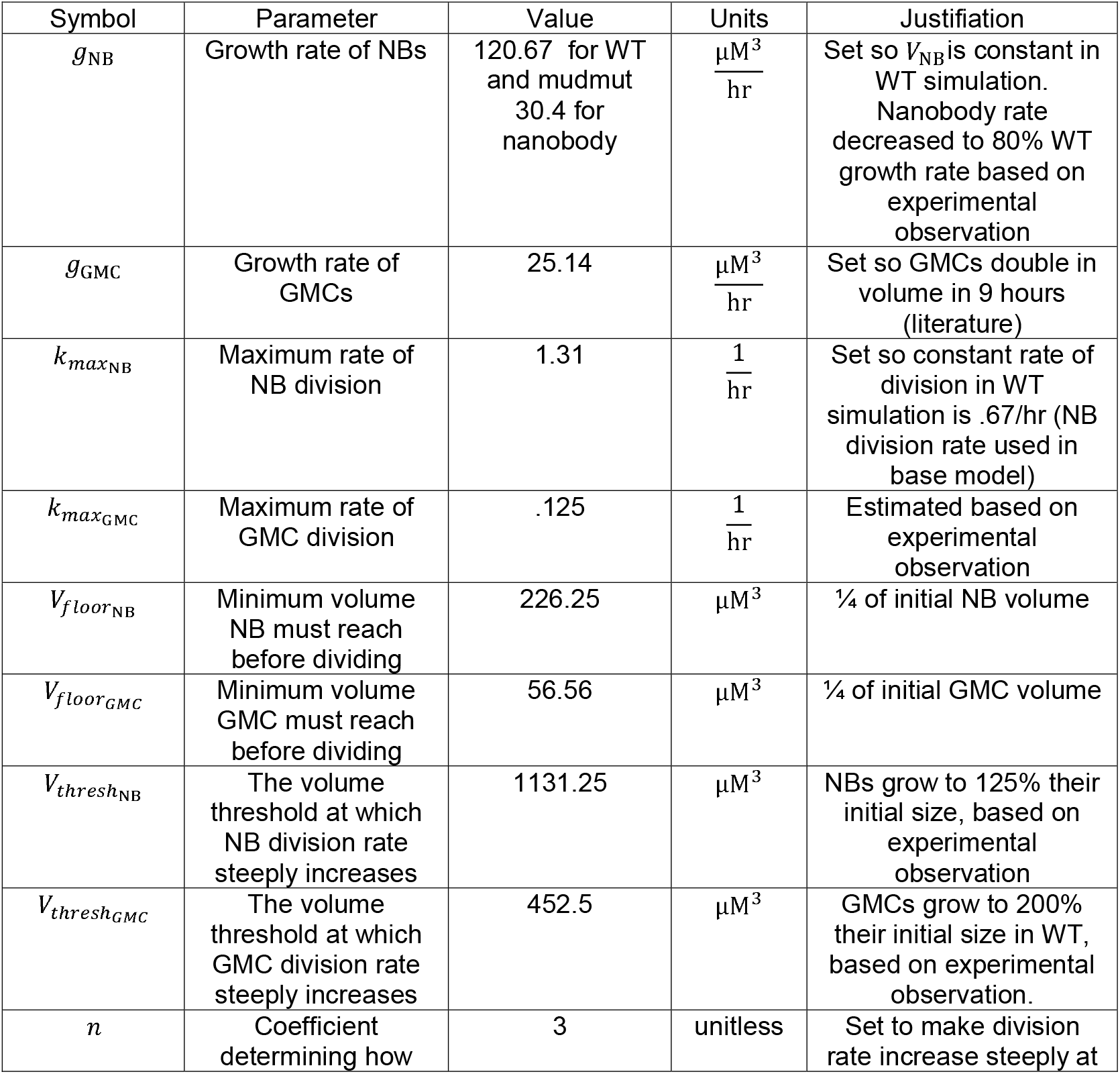
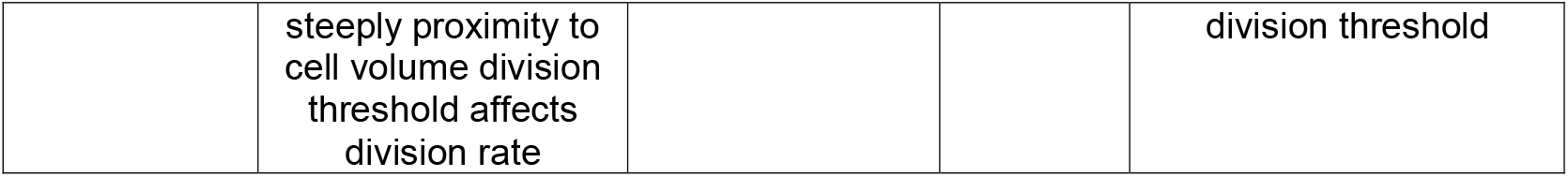

| Symbol | Parameter | Value | Units | Justification |
| --- | --- | --- | --- | --- |
| $g_{NB}$ | Growth rate of NBs | 120.67 for WT<br>and mudmut<br>30.4 for<br>nanobody | $\frac{\mu M^3}{hr}$ | Set so $V_{NB}$ is constant in<br>WT simulation.<br>Nanobody rate<br>decreased to 80% WT<br>growth rate based on<br>experimental<br>observation |
| $g_{GMC}$ | Growth rate of<br>GMCs | 25.14 | $\frac{\mu M^3}{hr}$ | Set so GMCs double in<br>volume in 9 hours<br>(literature) |
| $k_{max_{NB}}$ | Maximum rate of<br>NB division | 1.31 | $\frac{1}{hr}$ | Set so constant rate of<br>division in WT<br>simulation is .67/hr (NB<br>division rate used in<br>base model) |
| $k_{max_{GMC}}$ | Maximum rate of<br>GMC division | .125 | $\frac{1}{hr}$ | Estimated based on<br>experimental<br>observation |
| $V_{floor_{NB}}$ | Minimum volume<br>NB must reach<br>before dividing | 226.25 | $\mu M^3$ | ¼ of initial NB volume |
| $V_{floor_{GMC}}$ | Minimum volume<br>GMC must reach<br>before dividing | 56.56 | $\mu M^3$ | ¼ of initial GMC volume |
| $V_{thresh_{NB}}$ | The volume<br>threshold at which<br>NB division rate<br>steeply increases | 1131.25 | $\mu M^3$ | NBs grow to 125% their<br>initial size, based on<br>experimental<br>observation |
| $V_{thresh_{GMC}}$ | The volume<br>threshold at which<br>GMC division rate<br>steeply increases | 452.5 | $\mu M^3$ | GMCs grow to 200%<br>their initial size in WT,<br>based on experimental<br>observation. |
| $n$ | Coefficient<br>determining how | 3 | unitless | Set to make division<br>rate increase steeply at |
|  | steeply proximity to cell volume division threshold affects division rate |  |  | division threshold |

#### Model 2

Model 2 retains all of the ODEs from Model 1 and adds an additional state variable to track the changes in NB division threshold over time.

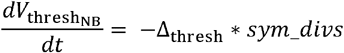

Where Δ_thresh_ is the constant amount the division threshold decreases with each symmetric division, and *sym*_*divs* is the number of symmetric divisions that have occurred in the simulation, equal to 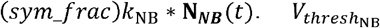 is initialized to 1131.25 µM^3^.

The new parameters are as follows.

| Symbol | Parameter | Value | Units | Justification |
| --- | --- | --- | --- | --- |
| $\Delta_{\text{thresh}}$ | Constant amount $V_{\text{thresh}_{\text{NB}}}$ is decreased with each symmetric division | 33.94 | $\mu\text{M}^3$ | 3% the initial division threshold. Calibrated so NBs do not reach their minimum volumes by the end of 48 hours in the simulation. |

#### Model 3

Model 3 takes Model 1 but determines *g*_NB_ using the following formula:

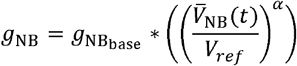

Where 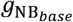 is the base NB growth rate, *V*_*ref*_ is a reference NB volume, and *α* is the sensitivity of the growth rate to the NB volume.

The new parameter values are as follows:

| Symbol | Parameter | Value | Units | Justification |
| --- | --- | --- | --- | --- |
| $g_{NB_{base}}$ | Base NB growth rate | 120.67 for WT and mudmut<br>96.53 for nanobody | $\mu M^3$ | Set to the growth rate in previous models |
| $V_{ref}$ | Volume at which $g_{NB} = g_{NB_{base}}$ | 905 | $\mu M^3$ | Set to the initial NB volume |
| $\alpha$ | Sensitivity of growth rate to NB volume | [1,5] | unitless | Capability of model to constrain mutant growth depends on this value |

#### Model 4

Model 4 takes Model 2 and makes *g*_NB_ dynamic as described in Model 3. All parameter values are as defined in Model 2 and Model 3.

#### Model 5

Model 5 builds on the Base Model by using the equation below to calculate *k*_NB_

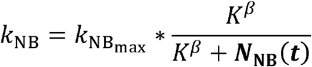

Where 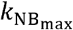 is the maximum NB division rate, K is the quantity of NBs at which the NB self-repression is half-maximal, and *β* is the Hill coefficient.

| Symbol | Parameter | Value | Units | Justification |
| --- | --- | --- | --- | --- |
| $k_{NB_{max}}$ | Maximum NB division rate | 1.30 | $\frac{1}{hr}$ | Set so the rate of division in WT simulations is constant |
| $K$ | The quantity of NBs at which NB self-repression is half-maximal | 2 | NBs | Set so repression is strong following the first symmetric division |
| $\beta$ | Sensitivity of growth rate to NB quantity | [1,5] | unitless | Capability of model to constrain mutant growth depends<br>on this value |

## Acknowledgements

We thank Melissa Harrison for the anti-Grh and Chris Doe for the anti-Hey antibody. We also thank Federico Tenedini with help with the RNAseq experiments and members of the Cabernard laboratory for helpful discussions and comments. This work was supported by the American Cancer Society (PF-24-1308526-01-CCB to J.I.H), the Washington Research Foundation (WRF to N. B) and the National Institutes of Health (R35GM148160 to C.C; NINDS R01 NS076614 to J.P).

## Supplemental Figure legends

**Figure S1: Nanobody-expressing neuroblasts divide asymmetrically, symmetrically or with inverted asymmetry**.

**(A)** Representative cartoon of the nanobody approach: the apically tethered, anti-GFP nanobody is equipped with a kinase domain from Rho-kinase, phosphorylating and constitutively activating Myosin’s regulatory subunit Spaghetti Squash (Sqh) upon binding of GFP to the vhhGFP. Consequently, activated Myosin will be retained on the apical neuroblast cortex ^29^. **(B)** Representative image sequences of wild-type and nanobody-expressing neuroblasts. All neuroblasts co-express UAS-mCherry::Jupiter (white) with worGal4 and endogenously tagged Sqh::EGFP (green). Representative symmetric and inverted asymmetric divisions are shown for nanobody-expressing neuroblasts. **(C)** Quantification of division modes for wild type, control, and nanobody-expressing neuroblasts. Each dot represents one ratio measurement. Average and standard deviation are shown overlaid with a scatter plot. Significance was tested with Bonferroni’s multiple comparisons test. Wild type vs control: p >0.9999. Wild type vs nanobody: p<0.0001. **(D)** Wild-type neuroblasts align the mitotic spindle in parallel to the intrinsic apical (blue) – basal (red) polarity axis, repeatedly generating large self-renewed neuroblasts (green) and small differentiating GMCs (red), dividing only once more. Trapping activated Myosin on the apical neuroblast cortex can induce **(E)** inverted asymmetric or **(F)** symmetric cell divisions with variable lineage trees. The resulting brains could contain neuroblasts of various sizes, many of which are smaller than normal wild-type neuroblasts. **(G)** Symmetric neuroblast divisions induced through spindle misalignment generate lineage trees with various outcomes, resulting in brains with potentially differently sized neuroblasts.

**Figure S2: Clusters 15 and 19 represent neuroblasts**

**(A)** Neuroblast clusters 15 and 19 highlighted on a UMAP plot. **(B)** DotPlot of marker genes used in the annotation of the neuroblast clusters.

**Figure S3: Cluster 11 represents INPs, and clusters 5, 7, 9, 12 represent GMCs, respectively**

**(A)** INP cluster 11 highlighted on a UMAP plot. **(B)** DotPlot of marker genes used in the annotation of the INP cluster. **(C)** GMC clusters 5, 7, 9, and 12 are highlighted on a UMAP plot. **(D)** DotPlot of marker genes used in the annotation of the GMC clusters.

**Figure S4: Immature, developing, and mature neurons separate into specific clusters**

**(A)** UMAP plot showing Newborn neuron (clusters 3, 13, 14) and immature neuron clusters (clusters 1, 2, 4, 8, 10, 18). **(B)** DotPlot of marker genes used in the annotation of the newborn neuron and immature neuron clusters. **(C)** UMAP plot showing developing and mature neurons. **(D)** DotPlot of marker genes used to annotate developing and mature neurons.

**Figure S5: Cluster 21 and cluster 22 represent Iro-C and Kenyon cells, respectively**

**(A, C)** UMAP plot highlighting cluster 21 containing mature Iro-C neurons and Kenyon cells, respectively. **(B, D)** DotPlot of marker genes used to annotate Iro-C neurons and Kenyon cells, respectively.

**Figure S6: New cluster markers and a comparison of immature and mature neuron numbers between genotypes**

**(A)** The top three differentially expressed genes per cluster (by log2FC) are shown on a DotPlot (see methods). Validated markers are not shown here. Red boxes outline the dots corresponding to the three genes per cluster for easier viewing. These are genes that have not been validated *in vivo* as cell type specific but are among the most significant principal components that distinguish clusters from one another and could be used as markers for cell typing in future RNA sequencing experiments. Immature **(B)** and mature **(E)** neuron clusters highlighted on UMAP plots for each condition, respectively. The total number of cells belonging to **(C)** immature neuron and **(F)** mature neuron clusters, normalized for library size and control. Immature neurons **(D)** and mature neurons **(G)** per neuroblast. Cells within immature neuron clusters with at least 1 *elav* transcript were used for the quantification for each condition. Cells within mature neuron clusters with at least 1 *nSyb* or *brp* transcript were used for the quantification for each condition.

**Figure S07: Model dynamics details**

Base model shown for **(A)** wild type and **(B)** *mud* mutant neuroblasts. The base model predicts that *mud* mutant NBs generate far more cells than wild-type NBs, inconsistent with experimental results.

**Figure S08: Mutant NB cell cycle durations continue increasing over time for models 3, 4, and 5**.

**(A)** NB Cell cycle duration is constant in the base model. **(B)** In model 1, volume activation increases cell-cycle duration to a new steady state in *mud* and nanobody-expressing NBs. **(C)** In model 2, a symmetric division threshold ultimately causes cell cycle duration to decrease to its minimum. Model 3 **(D)**, model 4 **(E)**, and model 5 **(F)** show the increase in cell cycle for mud mutant and nanodoby-expressing NBs. Models are only shown with an *α* = 3 (D, E) and β = 4 (F).

**Figure S9: Symmetrically dividing *mud***^***4***^ **mutant neuroblasts show asymmetric lineage behavior and extended interphase times**

**(A)** Schematic lineage tree of a symmetrically dividing neuroblast, illustrating the used nomenclature and lineage structure. Parental (P) neuroblasts generate an A and a B neuroblast after a symmetric division. A and B neuroblasts can either divide symmetrically or asymmetrically. Only dividing cells propagate the lineage and receive a lineage label. **(B)** Plot showing cell diameter for multiple generations in symmetrically dividing *mud*^*4*^ mutant neuroblasts. A and B neuroblasts are consistently smaller and do not regrow to parental NB size. A lineages are consistently longer than B lineages. **(C)** Cell cycle length comparison between wild type and *mud*^*4*^ mutant neuroblasts. Statistical test used: unpaired T-test. ** p < 0.01

**Figure S10: NB growth rate-sensitivity to NB volume (.) determines whether volume-sensitive growth effectively suppresses mutant NB proliferation when division threshold volume is constant**.

Cell counts shown for Model 3 run with different values of *α*.

**Figure S11: NB growth rate-sensitivity to NB volume (*α*) determines whether volume-sensitive growth effectively suppresses mutant NB proliferation when division threshold**

**volume is regulated by symmetric division**.

Cell counts shown for Model 4 run with different values of *α*.

**Figure S12: NB growth rate sensitivity to NB count (*β*) determines whether NB self-inhibition effectively suppresses mutant NB proliferation**. Cell counts shown for Model 5 run with different values of *β*.

**Figure S13: GO-terms for *mud* mutant and nanobody-expressig neuroblasts of differentially expressed genes**

GO term analysis for differentially expressed genes comparing **(A)** *mud*^*4*^ versus control, **(B)** nanobody-expressing neuroblasts versus control, and **(C)** genes differentially expressed in both *mud*^*4*^ and nanobody-expressing neuroblasts versus control. The combined score consists of a combination of the p-value and z-score. The z-score measures the deviation from the expected rank of the GO term. Dot size reflects number of associated genes and dot color the corresponding p-value. Genes are listed for each GO-term.

**Figure S14: Genes differentially expressed between wild type, *mud***^***4***,^ **and nanobody-expressing brains associate with neuroblast, differentiation, cell cycle, neurogenesis, and cytoskeleton functions**

Genes differentially expressed in **(A)** *mud*^*4*^, **(B)** nanobody-expressing, or **(C)** both *mud*^*4*^ and nanobody-expressing neuroblasts, compared to wild type, are associated with neuroblast, differentiation, cell cycle, neurogenesis, and cytoskeleton functions. Blue rectangles indicate keywords associated with the gene on the x-axis. Keywords are derived from GO terms. Pros ^42^ **(D)** and **(E)** Dpn ^103^ target genes that are differentially expressed in *mud*^*4*^ and nanobody-expressing brains. Green and red boxes indicate whether the genes are up- or downregulated in *mud*^*4*^ and nanobody-expressing neuroblasts compared to wild-type NBs.

**Figure S15: NB growth genes are not significantly differentially expressed in *mud***^***4***^ **or nanobody-expressing neuroblasts**

Volcano plots of DEGs between wild-type and *mud*^*4*^ (left), and wild-type and nanobody (right), highlighting genes in the **(A)** Insulin pathway ^95^, **(B)** the Mediator complex ^50^, **(C)** the oxidative phosphorylation pathway ^50^, or **(D)** other genes regulating NB growth ^96,97^. We only considered genes that are expressed in at least 25% of neuroblasts, with a p-value of 0.001 or better, and a minimum log2-fold change of 0.5. Any gene appearing in one plot but not the other signifies that that gene was expressed by less than 25% of cells in that group.

## Movie legends

**Movie 1: Wild type neuroblasts divide asymmetrically**

Representable wild type neuroblast expressing Sqh::EGFP (cortex; green) and the mitotic spindle marker mCherry::Jupiter (white; driven by worGal4). The corresponding lineage tree is shown at the end of the movie. Scale bar is 5μm. Time is in hours:minutes:seconds.

**Movie 2: Nanobody-expressing neuroblasts can divide in an inverted asymmetric fashion** Representable Nanobody-expressing neuroblast co-expressing Sqh::EGFP (cortex; green) and the mitotic spindle marker mCherry::Jupiter (white; driven by worGal4). The first division generates a small apical and a large basal cell. The large basal cell divides again at 4:05:00. The corresponding lineage tree is shown at the end of the movie. Scale bar is 5μm. Time is in hours:minutes:seconds.

**Movie 3: Nanobody-expressing neuroblasts can divide symmetrically by size**. Representable Nanobody-expressing neuroblast co-expressing Sqh::EGFP (cortex; green) and the mitotic spindle marker mCherry::Jupiter (white; driven by worGal4). The first division generates two equal sized cells. The apical cell continues to divide symmetrically two more times (at 2:57:44 and 7:34:17) before the basal cell divides (at 13:44:42). The corresponding lineage tree is shown at the end of the movie. Scale bar is 5μm. Time is in hours:minutes:seconds.

**Movie 4: Wild type neuroblasts retain Dpn at higher levels than differentiating GMCs**.

Wild-type neuroblast expressing Sqh::GFP (shown in green) and tdTomato::Dpn (shown in white). Green dotted line at 0:00:00 outlines the neuroblast of interest with strong nuclear Dpn signal. At 1:09:00, the neuroblast enters mitosis, releasing nuclear Dpn into the cytoplasm. At 01:24:00, the cell divides asymmetrically and Dpn signal reappears in the nuclei of both newly formed neuroblast (green dotted line) and the differentiating GMC (red dotted line). The neuroblast’s signal is brighter in the self-renewed neuroblast (apical cell). Over time, Dpn signal fades away in the GMC, while being retained in the self-renewed neuroblast. Scale bar is 5μm. Time is in hours:minutes:seconds.

**Movie 5: In symmetrically dividing *mud***^***4***^ **mutant neuroblasts. Both sibling cells retain nuclear Dpn**.

Symmetrically dividing *mud*^*4*^ mutant neuroblast expressing Sqh::GFP (shown in green) and tdTomato::Dpn (shown in white). The neuroblast of interest (green dotted line) contains strong nuclear Dpn signal. At 0:27:00 the neuroblast enters mitosis, releasing Dpn into the cytoplasm. At 0:42:00, the NB divides symmetrically with equal amounts of Dpn reappearing in both sibling cell nuclei (Green and red dotted lines as show in in Figure 2C). Dpn signal remains roughly equal in the two daughter cells until the end of the movie. Scale bar is 5μm. Time is in hours:minutes:seconds.

**Movie 6: In symmetrically dividing nanobody-expressing neuroblasts both siblings retain nuclear Dpn**.

Nanobody-expressing neuroblast, coexpressing Sqh::GFP (shown in green) and tdTomato::Dpn (shown in white). The neuroblast of interest is highlighted with the green dotted line at 0:00:00. At 0:24:00, the neuroblast enters mitosis, releasing Dpn into the cytoplasm. At 0:39:00, the NB divides symmetrically and Dpn signal reappears in the nuclei of both newly formed daughter cells with roughly equal fluorescence intensity (highlighted with green and red dotted lines as shown in Figure 2E). Despite bleaching, both daughter cells contain roughly equal tdTomato::Dpn. Scale bar is 5μm. Time is in hours:minutes:seconds.

**Movie 7: Nanobody-expressing neuroblasts can generate a small apical Dpn**^**-**^ **and larger basal Dpn**^**+**^ **siblings**.

Nanobody-expressing neuroblast, coexpressing Sqh::GFP (shown in green) and tdTomato::Dpn (shown in white). The neuroblast of interest is highlighted with the green dotted line at 1:54:00. The neuroblast divides with inverted asymmetry, generating a small apical (green dotted line) and large basal cell (red dotted line) at 2:42:00m whereby only the larger basal cell contains tdTomato::Dpn. Dpn signal remains high in the basal cell but is absent from the apical cell. Scale bar is 5μm. Time is in hours:minutes:seconds.

**Movie 8: Nanobody-expressing neuroblasts can generate a small apical Dpn**^**+**^ **and larger basal Dpn**^**+**^ **siblings**.

Nanobody-expressing neuroblast, coexpressing Sqh::GFP (shown in green) and tdTomato::Dpn (shown in white). The neuroblast of interest is highlighted with the green dotted line at 0:18:00. The neuroblast divides with inverted asymmetry, generating a small apical (green dotted line) and large basal cell (red dotted line) at 2:18:00. Both sibling cells show nuclear tdTomato::Dpn. Scale bar is 5μm. Time is in hours:minutes:seconds.

